# Regulatory mutants of the *Tbx1* gene alter transcription programs of lineage determination and patterning in early mesoderm

**DOI:** 10.64898/2026.08.08.743664

**Authors:** S Allegretti, O Lanzetta, M Bilio, R Ferrentino, P Salerno, P Zoppoli, G Merla, C Angelini, A Baldini

## Abstract

The *Tbx1* gene is haploinsufficient in mice and in humans, where it causes a DiGeorge syndrome phenotype characterized by developmental deficits of the pharyngeal apparatus. TBX1 plays a critical role in the differentiation and regionalization of the cardiopharyngeal mesoderm lineage and its derivatives. Nevertheless, its regulation is incompletely understood. Here we used a combination of computational and wet-lab approaches to identify regulatory sequences of the *Tbx1* gene, and we use single-cell molecular analysis as a read-out and to establish the consequences of their deletion. Results revealed a cluster of regulatory sequences with at least three distinct elements. Elimination of the entire cluster caused a near shut down of the gene, while individual deletions had milder, quantitative effects. Transcriptomic analyses of the deletion mutants revealed the down regulation of genes related to cardiopharyngeal lineage specification and, more surprisingly, up regulation and anteriorization of genes related to embryonic patterning, thereby providing a rationale for the severe dysmorphogenesis of the posterior pharyngeal apparatus observed in *Tbx1* mutant mice.

## INTRODUCTION

Research into the regulation of *Tbx1* gene expression has been relatively sparse despite the development importance of the gene. *Tbx1* encodes a T-box transcription factor that is essential for pharyngeal apparatus and cardiac outflow tract (OFT) development, and for the formation of craniofacial structures (Baldini et al. 2017; Jerome and Papaioannou 2001; Lindsay et al. 2001; Merscher et al. 2001). In vertebrates, a fundamental contribution to the development of these structures is provided by the cardiopharyngeal mesoderm (CPM), an evolutionarily conserved multipotent cell lineage that gives rise to second heart field (SHF) and branchiomeric muscle progenitors (Argiro et al. 2024; Devine et al. 2014; Diogo et al. 2015; Lescroart et al. 2014, 2018, 2022). Among the transcription factors that drive these multipotent progenitors through morphogenetic development, *Tbx1* plays a central role. In the mouse, *Tbx1* is expressed from embryonic stage E7.5 in the pharyngeal endoderm and ectoderm, mesodermal cores of the pharyngeal arches, paraxial mesoderm, and SHF cells. In this context, *Tbx1* is essential for the proliferation, survival, and timely deployment of SHF progenitors to the OFT, right ventricle, and to portions of the atria (Huynh et al. 2007; Rana et al. 2014; Xu et al. 2005). *Tbx1* also maintains the extracellular matrix–cell interactions that are required for OFT morphogenesis (Alfano et al. 2019). These functions make *Tbx1* a key regulator of progenitor maintenance within the CPM, thereby preventing premature differentiation and guaranteeing proper integration of cardiac and craniofacial developmental programs. The importance of this developmental system, with clinical relevance, is highlighted by the DiGeorge syndrome phenotype, part of the 22q11.2 deletion syndrome spectrum, in which *Tbx1* haploinsufficiency is the primary genetic determinant. Despite its central role in development and disease, the regulatory architecture controlling *Tbx1* gene expression remains only partially defined. Previous studies identified a 200 bp sonic hedgehog (Shh) responsive enhancer upstream of *Tbx1* (−12.8 kb from TSS) containing FOX transcription factor binding sites, suggesting that SHH signalling maintains *Tbx1* expression through FOXA2, FOXC1, and FOXC2 in pharyngeal tissues (Brown et al. 2004; Garg et al. 2001; Hu et al. 2004; Maeda et al. 2006; Yamagishi et al. 2003). Subsequent mutagenesis and transgenic analyses revealed multiple conserved noncoding sequences, including a pharyngeal endoderm specific enhancer and an intronic suppressor element, suggesting the existence of a multiple element repertoire that contributes to *Tbx1* gene regulation (Zhang 2010). This architecture is reminiscent to the enhancer structure controlling other developmental genes, such as *Shh*, the expression of which depends upon long range regulatory domains spanning 900–1000 kb of DNA composed of multiple tissue specific enhancers to achieve precise spatiotemporal expression (Anderson and Hill 2014).

The combination of computational methods with advanced cellular models, gene editing, and single-cell transcriptomics and epigenomics offers new possibilities for redefining the strategies used to identify and validate regulatory sequences. The CRISPR-Cas9 tool makes it relatively easy to manipulate multiple specific genomic regions in mammalian cells (Cong et al. 2013; Jinek et al. 2012). Furthermore, advances in sequencing technologies allow the rapid identification of candidate regulatory elements. In particular, simultaneous chromatin accessibility and RNA assays, has opened the door to powerful new strategies to the identification of putative regulatory sequences, providing a direct link between the state of chromatin in specific genomic regions and the expression level of their target genes (Preissl et al. 2023; Racioppi et al. 2019; Ryan and Farley 2020).

Here we used multiomic datasets to identify putative regulatory sequences with the help of a machine learning procedure (Lanzetta et al. 2025). Through this, we identified an 8.5Kbp cluster of regulatory sequences upstream of the mouse *Tbx1* gene that includes multiple regulatory elements. Gene editing, single nucleus RNA-seq and extensive transcriptomic analyses provided evidence of complex regulatory logic and identified novel pathways affected by gene dysregulation.

## RESULTS

### Identification of high scoring putative regulatory sequences (PRSs) in the Tbx1 genomic locus

We used a multiomic dataset obtained from a cardioid differentiation model to study single nucleus (sn) ATAC-seq peaks in a 40kbp genomic window of the mouse *Tbx1* locus (Fig. 1a). Within this segment, the peak calling software identified 14 ATAC-seq peaks located upstream, within, and downstream of the gene. To this set of peaks, we applied a previously described machine-learning strategy trained on known enhancers (Aurigemma et al. 2024) to score the peak sequences for their probability to be enhancers. Results shown in Tab. 1 indicate the values for each of the sequences, expressed in a range from 0 (0% probability to be enhancers) to 1 (100% probability), we considered positive a score of >0.5. Visual inspection of the ATAC coverage map divided by cell cluster (Fig. 1b) identified a 7.6Kbp region that included 3 variable ATAC peaks (named PRS10, PRS11, and PRS12) located approx. 20 kb upstream to the *Tbx1* transcription start site (TSS). Hereafter, we refer to this region as a cluster of regulatory elements or CLRE. PRS10 and PRS12 had a positive prediction score (0.67 and 0.65, respectively), while PRS11 had a low score (0.33). We next tested whether these peaks were present in mouse embryo tissues of a developmentally relevant stage (E9.25) by reanalysing and querying a publicly available snATAC-seq dataset (Ranade et al. 2022); results showed that PRS10, 11, and 12 peaks were indeed detected in mouse embryo tissues (Suppl. Fig. 1). Next, we used available chromatin capture data (HiC, data.4dnucleome.org) to infer whether the CLRE is likely to interact with the *Tbx1* promoter region. Overlaying CLRE coordinates with HiC data revealed a topologically associating domain (TAD) that included the CLRE and the *Tbx1* 5’ region (Suppl. Fig. 2). Therefore, we focused on this CLRE for further studies.

**Figure 1.**
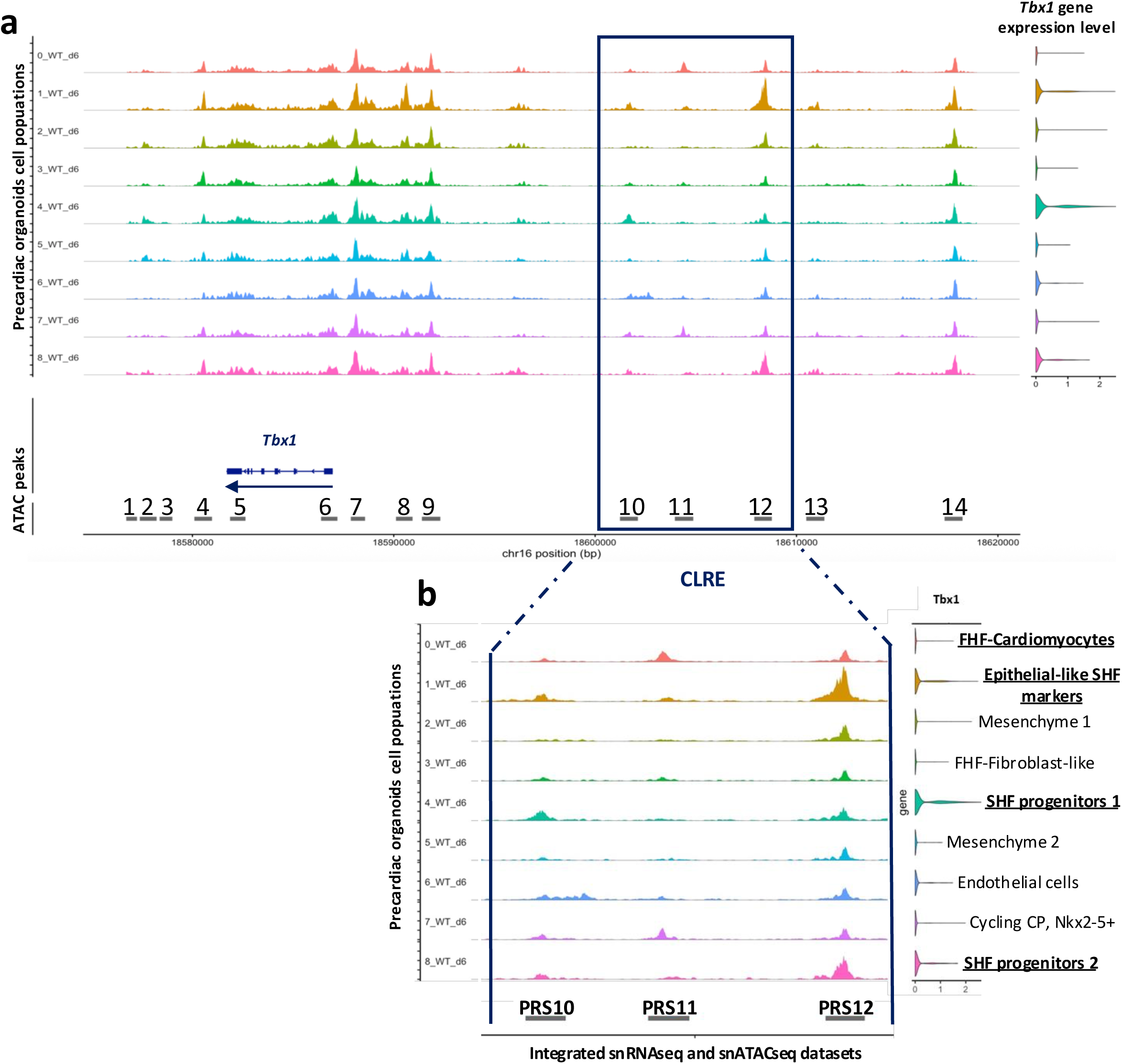
Identification of putative regulatory sequences (PRS) within the mouse Tbx1 gene locus. Coverage of snATAC-seq data in a 40kbp genomic window of the mouse *Tbx1* locus. We used a published multiomic dataset (simultaneous snATAC-seq and snRNA-seq) from precardiac organoids (Lanzetta et al. 2025). a) The genome browser window shows the 14 peaks identified in the dataset divided into 9 cell clusters of WT cells; *Tbx1* expression level for each cell cluster is shown on the right. b) The panel focuses on a region (cluster) of approx. 7.5 Kbp (CLRE) that includes the three peaks that we named PRS10, PRS11 and PRS12. The identity of the cell clusters, as previously reported (Lanzetta et al. 2025), is shown on the right. *Tbx1* expression is most prevalent in the clusters shown in bold.

**Table 1.** Probability scores of snATAC-seq/PRS regions of the *Tbx1* locus.

| Coordinates (mm10 assembly) | Average prediction score | PRS ID |
| --- | --- | --- |
| chr16-18576567-18577247 | 0.39 | 1 |
| chr16-18577384-18578222 | 0.37 | 2 |
| chr16-18578371-18579005 | 0.41 | 3 |
| chr16-18580095-18580974 | **0.53** | **4** |
| chr16-18581868-18582636 | 0.36 | 5 |
| chr16-18586388-18587203 | **0.64** | **6** |
| chr16-18587866-18588576 | 0.38 | 7 |
| chr16-18590110-18590935 | 0.12 | 8 |
| chr16-18591397-18592314 | 0.43 | 9 |
| chr16-18601232-18602126 | **0.68** | **10** |
| chr16-18603959-18604876 | 0.33 | 11 |
| chr16-18607904-18608777 | **0.65** | **12** |
| chr16-18610483-18611385 | 0.32 | 13 |
| chr16-18617351-18618243 | 0.37 | 14 |

Positive prediction scores are shown in bold.

We noted that PRS10 also included a TBX1 binding site previously identified in mouse embryo material by ChIP-seq (Nomaru et al., 2021). Therefore, we checked whether the ATAC peak amplitude (coverage) was affected by the loss of *Tbx1* function. Visual inspection showed that in *Tbx1*^-/-^ cells, the snATAC-seq coverage was lower compared to wildtype (WT), particularly in clusters c2 and c4,identified as two subpopulations of anterior CPM and somatic mesoderm in the original publication (Lanzetta et al. 2025) (Fig. 2). Analogously, we found reduced coverage in two clusters of a published dataset of snATAC-seq in *Tbx1*^-/-^ vs WT mouse embryos at E9.25 (Ranade et al. 2022) (Suppl. Fig. 3). In both cases, reduced coverage appeared only in the PRS10 region and only in selected clusters. Thus, *Tbx1* function contributes to chromatin opening of the PRS10 region, at least in selected contexts, suggesting positive feedback.

**Figure 2.**
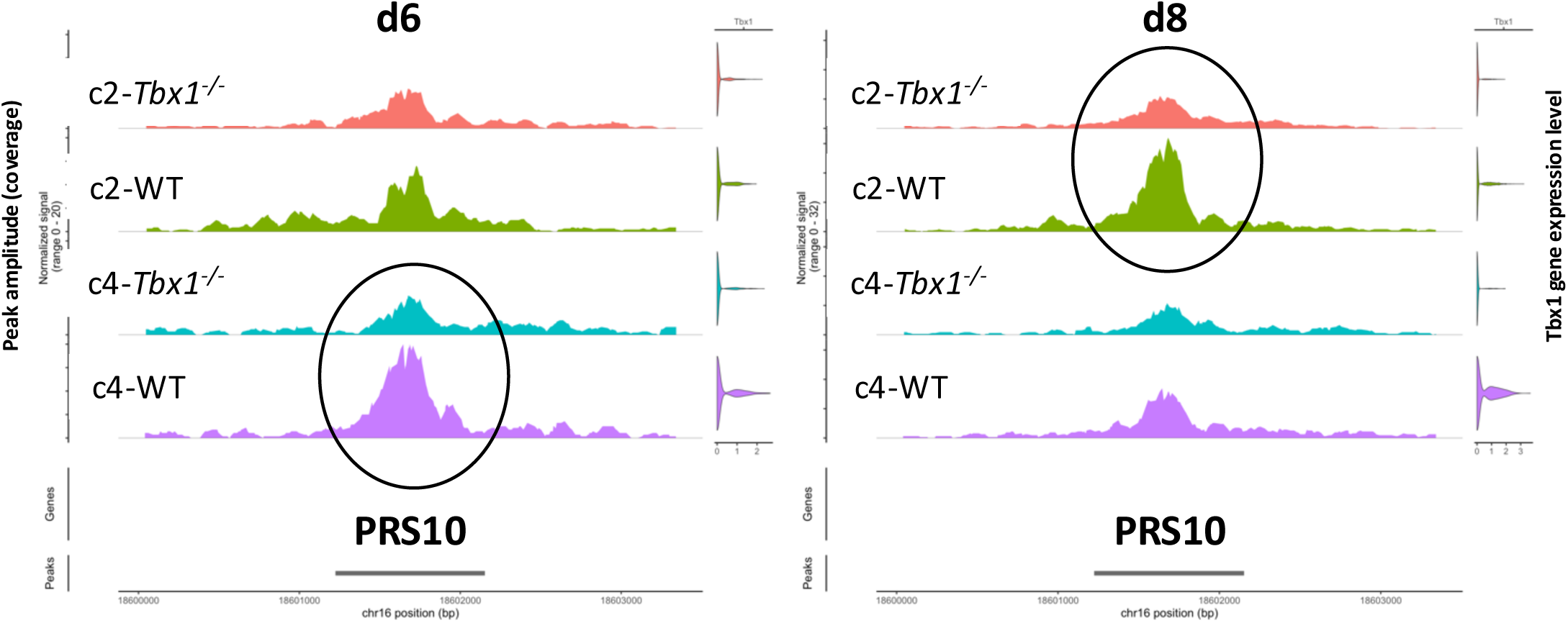
PRS10 accessibility responds to Tbx1 genotype in different clusters and timepoints. In the absence of *Tbx1*, PRS10 chromatin accessibility is reduced in specific cell populations (black circles) (c2 and c4) at two stages of differentiation (day 6 and 8), suggesting a positive feedback autoregulatory mechanism. C2 and c4 were identified as two subpopulations of anterior cardiopharyngeal mesoderm (CPM) and somatic mesoderm (Lanzetta et al. 2025).

### Generation of deletion mutants within the CLRE domain and validation of PRSs

To understand whether the CLRE is required for *Tbx1* gene expression, we homozygously deleted the entire 7.6 KBp region using CRISPR-Cas9 technology (Fig. 3a and Suppl. Fig. 4). We obtained two clones, that were confirmed by sequencing, and subjected them to a cell differentiation protocol (Andersen et al. 2018). We assayed *Tbx1* gene expression at three stages of differentiation (day 4, 6 and 8) using quantitative reverse transcription PCR (qRT-PCR). Results showed a significant, strong down regulation of *Tbx1* gene expression in deleted clones, as compared to the parental cell line (Fig. 3a-a’). Thus, the CLRE is required for *Tbx1* gene expression in this differentiation model. Next, we asked how the individual ATAC PRSs contribute to *Tbx1* gene expression. To this end, we generated mouse ESC clones with homozygous deletion of each of the three regions (PRS10, 11, and 12) using CRISPR-Cas9 (Suppl. Fig. 4 and 5). The sequences of RNA guides used in these experiments are listed in Suppl. Table 1. We subjected deleted, sequence-verified clones to *in vitro* differentiation and assayed them for *Tbx1* gene expression. Results showed severe reduction of *Tbx1* gene expression in both ΔPRS10 and ΔPRS12 compared to the WT parental line (Fig. 3b-c), while expression in ΔPRS11 cells was not significantly different from the parental line (Fig. 3d).

**Figure 3.**
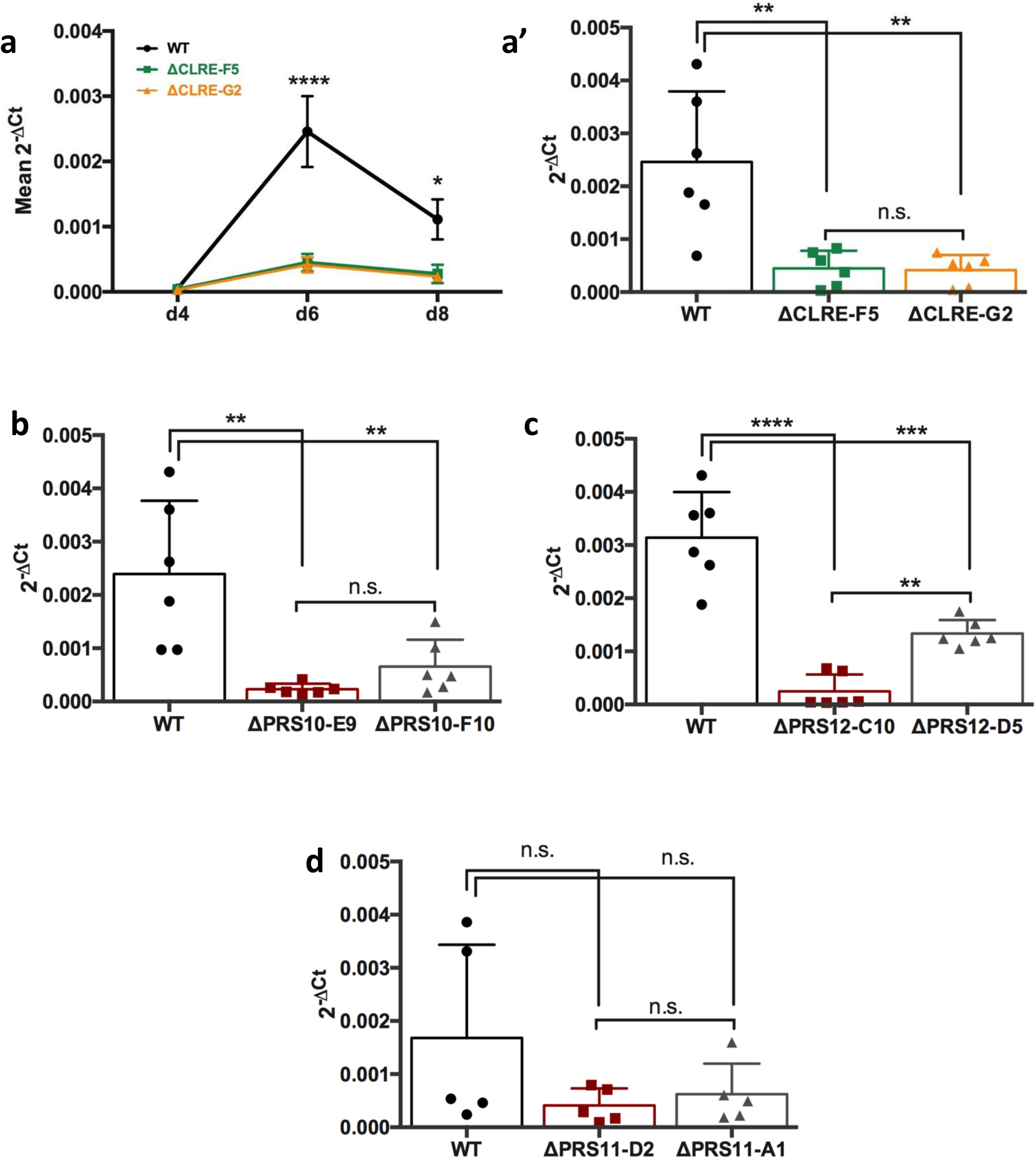
Deletion of CLRE, PRS10 and PRS12 but not PRS11 significantly reduced Tbx1 expression in precardiac organoids. a) Two independent clones carrying a homozygous deletion of the entire CLRE region and cells of the parental line (WT) tested for quantitative expression of *Tbx1* mRNA at days 4, 6, and 8 of differentiation; a’) same clones as in (a) in repeated experiments at d6. b) two independent clones carrying a deletion of PRS10 at d6; c) two independent clones with a deletion of PRS12 at d6; d) Two independent clones carrying the deletion of PRS11 at d6. Data from at least 5 independent differentiation experiments are presented as mean ± SEM (panel a) and ± SD (other panels) with individual data points shown. Two-way (panel a) and one-way (other panels) ANOVA with Tukey’s multiple comparisons test was applied for statistical analysis. Asterisks refer to adjusted P values: ****: P<0.0001; ***: P= 0.0001 ≤ p < 0.001; **: P= 0.001 ≤ p < 0.01; *: P= 0.01 ≤ p < 0.05; n.s. not significant.

### snRNA-seq data show diversity of regulatory elements functions

To gain information as to the relevance of our selected PRSs across cell types, we performed snRNA-seq of differentiated cells at day 6 from ΔPRS lines and the WT parental line, on two biological replicates of each cell line. We sequenced approx. 10000 nuclei per sample, for a total of approx. 80000 cells. After filtering for low quality cells and doublets (see Methods for details) we integrated the four genotypes and replicates and performed clusterization. Fig. 4a shows the Uniform Manifold Approximation and Projection (UMAP) display of the data. With the clustering resolution adopted, we distinguished 11 clusters, the top 20 marker genes of which are listed in Suppl. Tab. 2. Using comparison to published datasets (Tyser et al. 2021) and literature searches, we defined the clusters as shown in Tab. 2. *Tbx1* expression was mainly observed in clusters 2, 9, and 10 (Fig. 4b), identified as pharyngeal/splanchnic mesoderm, axial mesoderm, and endothelial cells, respectively. In Fig. 4c are shown the UMAPs divided by genotype and in Fig. 4d the relative *Tbx1* gene expression, which is visibly reduced in ΔPRS10 and 12. We pooled all cells from these three clusters into a single group and performed quantitative expression analyses comparing each genotype against WT using the Wilcoxon rank-sum test. Resulting differentially expressed genes (DEGs, P value adjusted <0.05) are listed in Suppl. Tab. 3. The Venn diagram shown in Fig. 5a illustrates the overlap between the DEGs observed in deleted mutants compared to the WT parental line. While there was a broad overlap between the three sets of DEGs, it was also evident that each genotype was associated with distinct transcriptomes. *Tbx1* expression was strongly down regulated in the ΔPRS10 line (avg. log2FC -4.3, adj *P*<<10^−6^) compared to WT cells, while in ΔPRS12 cells the down regulation was to an average log2FC -2.5 (adj *P*<<10^−6^). In ΔPRS11 cells, *Tbx1* expression was not significantly different from WT. This finding is consistent with qRT-PCR results obtained from the entire cell population in repeated differentiation experiments (Fig. 3d), but it is in apparent contrast with the partial overlap between ΔPRS11 vs. WT DEGs and ΔPRS10 (or ΔPRS12) vs. WT DEGs, which suggests a related transcriptional perturbation (Fig. 5a). For example, comparing DEGs in ΔPRS11 with DEGs in ΔPRS10, there is a large set of genes downregulated in both groups (Fig. 5b, bottom-left quadrant). We used multidimensional scaling of the full dataset to evaluate the Euclidian distance between the transcrptomes of all genotypes, the transcriptome of ΔPRS11 cells was closer to that of WT cells than ΔPRS10 and ΔPRS12 (Fig. 5c). This suggests that the deletion of PRS11 in this differentiation system has mild consequences compared to PRS10 and PRS12.

**Figure 4.**
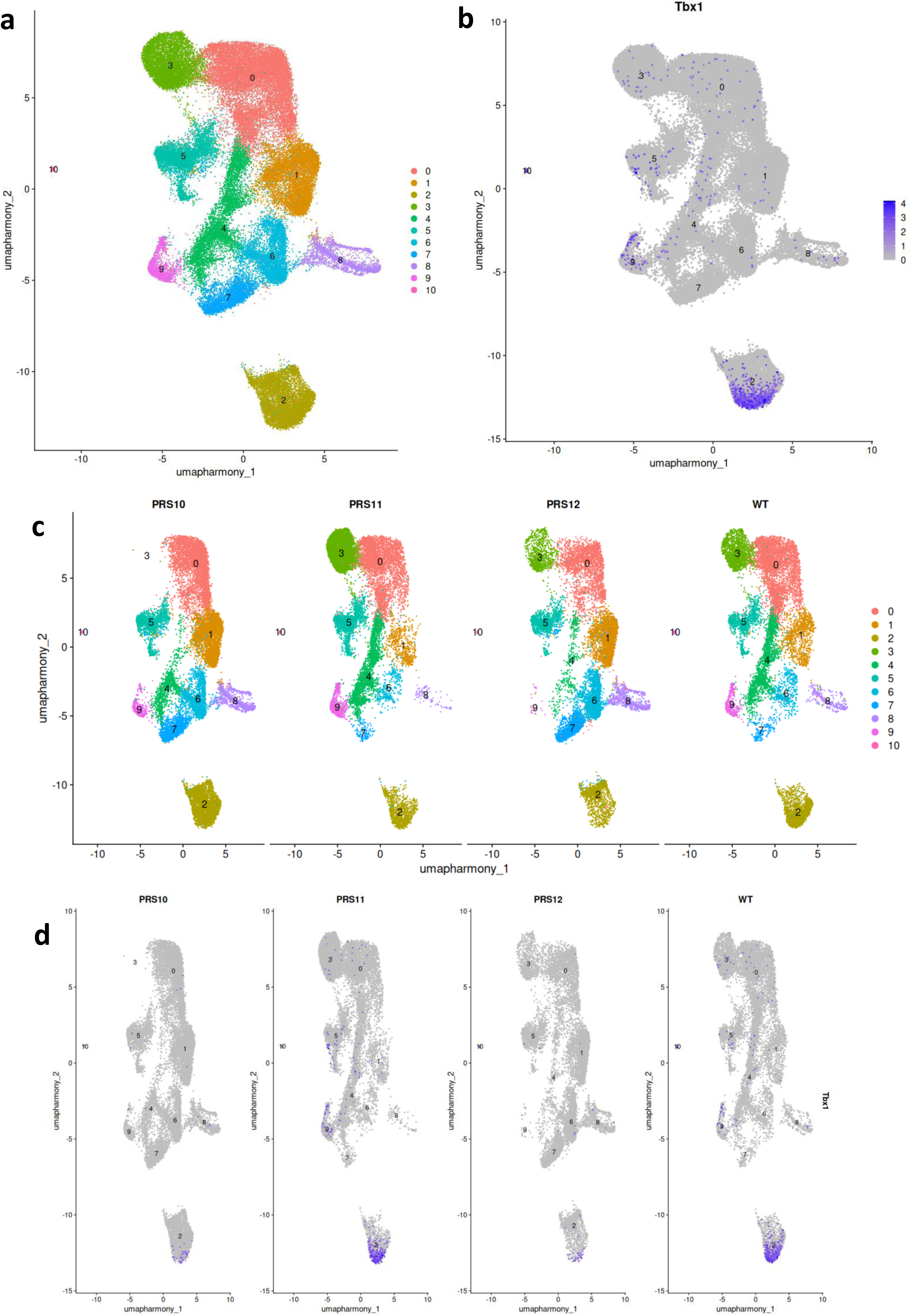
Tbx1 gene expression is severely affected by PRS deletions in specific subpopulations of differentiating organoids. a) UMAP representation of integrated snRNA-seq data from ΔPRS10, 11, 12 and parental cells. The population is divided into 11 clusters, the identification of which is listed in Table 2; b) *Tbx1* gene expression is mostly confined to clusters 2, 9, and 10; c) UMAP divided by genotype; d) *Tbx1* gene expression per each genotype: note similar expression between WT and ΔPRS11, but visibly reduced in ΔPRS10 and ΔPRS12, compared to WT cells. A residual population of cells expressing *Tbx1* is conserved in all mutants.

**Figure 5.**
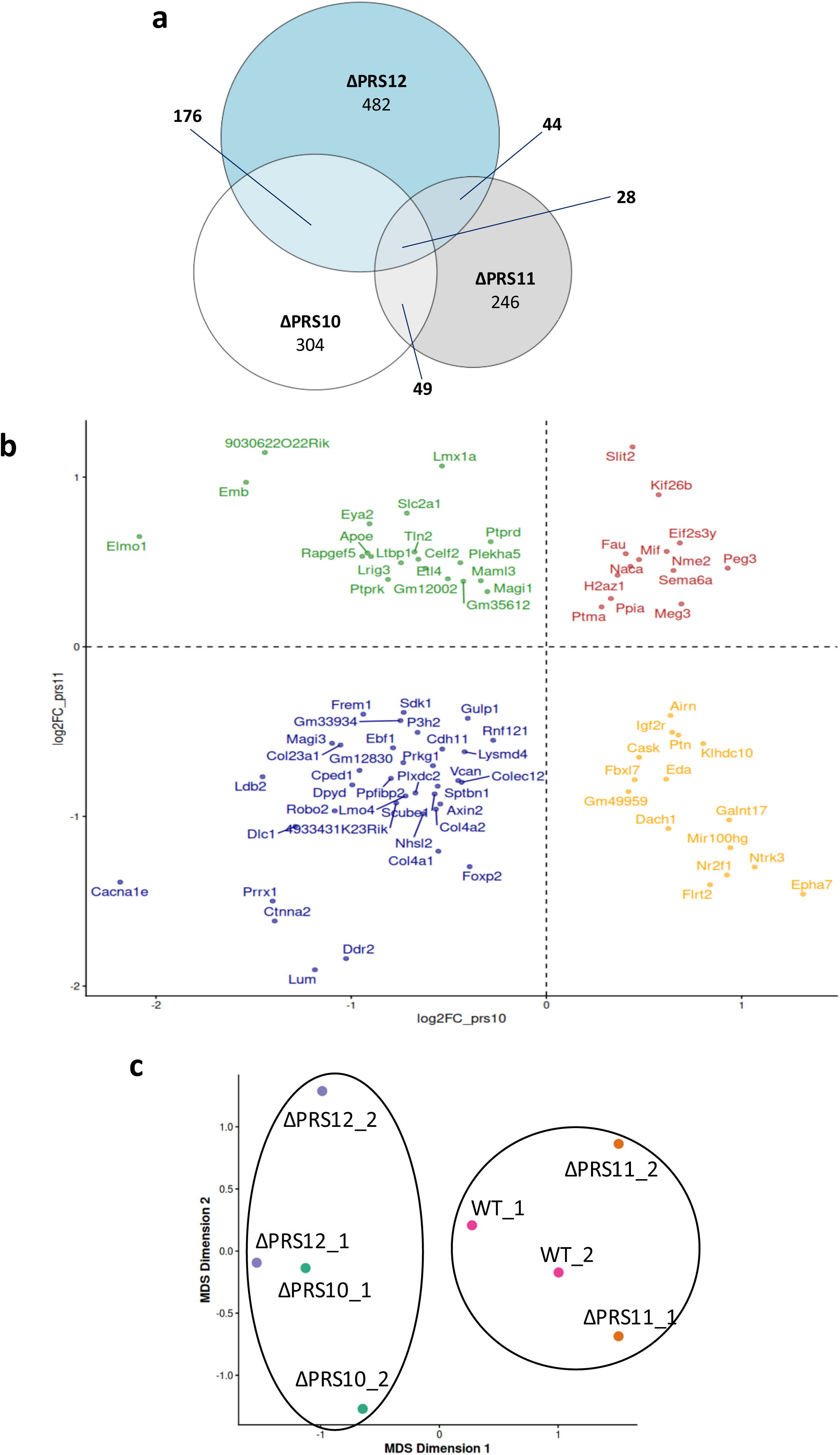
The regulatory mutants have distinct transcriptomes but exhibit partial overlap of differentially expressed genes, relative to WT cells. a) Venn diagram showing the overlap between differentially expressed genes (DEGs) in the three regulatory mutants relative to WT, down and up regulated genes are combined. b) The 4-quadrant diagram shows DEGs dysregulated in both ΔPRS10 and ΔPRS11, note that the largest subgroup is made of genes downregulated in both lines (bottom left quadrant). DEGs calculation was performed using only cells of clusters 2, 9, and 10 (Fig. 4a). c) The full transcriptomes of the three regulatory mutants and WT cells were subjected to multi-dimensional scaling calculation; each centroid represents a replicate of each genotype (two biological replicates per genotype). Note that WT and ΔPRS11 transcriptomes are closer with each other compared to ΔPRS10 and ΔPRS12.

**Table 2.** Cell type identification of full snRNA-seq full dataset.

| Cluster definition of integrated full dataset |  |
| --- | --- |
| Cluster | predicted ID |
| 0 | Neuroepithelial / early neural progenitor cells |
| 1 | Dorsal anterior neuroectoderm |
| **2** | **Pharyngeal/splanchnic mesoderm** |
| 3 | Anterior neural plate / early forebrain neuroepithelium |
| 4 | neuroectoderm |
| 5 | housekeeping genes, undefined |
| 6 | Dorsal Neural Tube / Early Neural Crest Progenitors |
| 7 | Hox-enriched module, A/P patterning, posterior trunk, neuromesoderm. |
| 8 | Pan-neuronal / maturing neuron signature |
| **9** | **Axial mesoderm** |
| **10** | **Endothelial cells, endothelial cell progenitors** |

Bold: Clusters exhibiting high *Tbx1* gene expression.

To reduce dilution from non-relevant cells, we focused our analyses to a subset of clusters enriched for *Tbx1*-expressing cells, i.e. clusters 2, 9, and 10 of Fig. 4a. We re-clustered this sub population and obtained 6 sub-clusters identified as shown in Fig. 6a-b (marker genes listed in Suppl. Tab. 4), hereafter indicated as S-0 to S-5. We used RNA velocity, implemented by *scvelo* (Bergen et al. 2020), to predict the trajectory relationship between the clusters. Results suggested that the mesenchymal transcriptome of cluster S-0 is likely to be the least differentiated cluster, progressing into early mesoderm progenitors (cluster S-1) and then into mesoderm with axial features (cluster S-2) (Fig. 6c). *Tbx1* expression was present in all the clusters in WT cells, except cluster S-0, but it was most prominent in cluster S-1 (Fig. 6d-e). In ΔPRS11 cells, the pattern of expression was similar to that of WT cells, but in ΔPRS10 and ΔPRS12 cells, the expression in cluster S-1 was strongly reduced, and it was very low or undetectable in cluster S-2 (Fig. 6d-e). Thus, the consequences of PRS10 and PRS12 deletions on the expression of *Tbx1* gene were evident in the overall population, although they were more pronounced in cluster S-2, which, according to RNA velocity assay, represents a cell population downstream (more differentiated) compared to cluster S-1, suggesting differentiation impairment.

**Figure 6.**
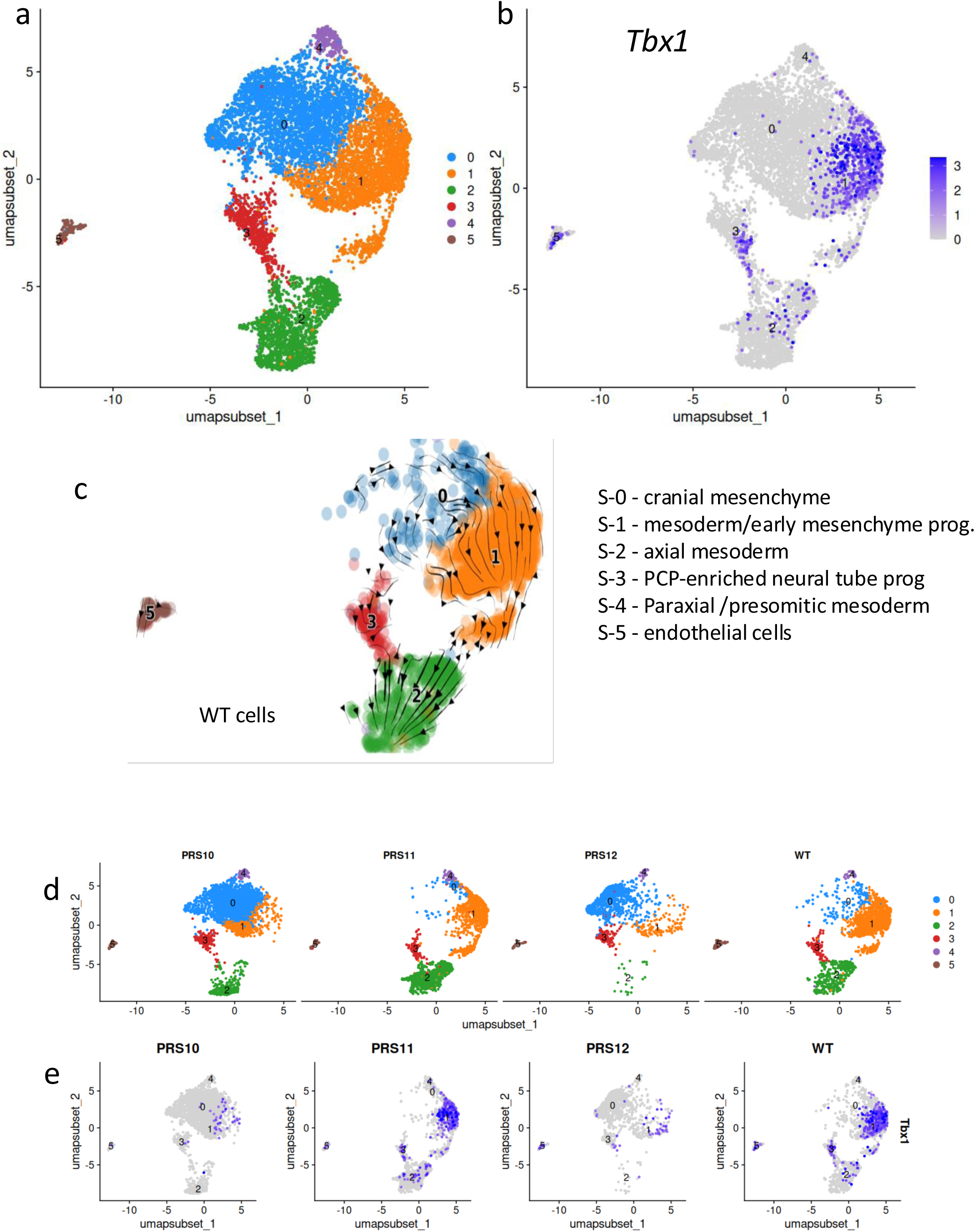
Inferring a differentiation trajectory within Tbx1-expressing early mesoderm. a) Integrated UMAP of cells from clusters c2, c9, and c10 (Fig. 4a) re-clustered into 6 subset-clusters; b) *Tbx1* gene expression in the subset is mostly restricted to S-1, S-2, and S-3; c) RNA velocity of WT cells suggests a trajectory 0 -> 1 -> 2 (early cranial mesenchyme -> mesoderm progenitors -> axial mesoderm) in this cellular model of differentiation. d) UMAP representation of the subset populations divided by genotype. Note that cluster S-0 is mostly present in ΔPRS10 and ΔPRS12 mutants; e) In ΔPRS10 and ΔPRS12 mutants, residual *Tbx1* expression is detected almost exclusively in cluster S-1.

### The transcriptional architecture of regulatory mutants highlights anomalies in gene networks involved in lineage determination and embryo patterning

We next explored additional transcriptional consequences of regulatory mutants by defining sets of co-regulated genes. For this, we used high dimensional weighted gene co-expression network analysis (hdWGCNA) (Childs et al. 2024; Morabito et al. 2023) to identify the transcriptional modules that were active in our cell differentiation model. We performed the analysis on WT cells of the subset described above to avoid interference from mutation-induced transcriptional changes. Results revealed 13 modules grouped in four categories (Fig. 7a, Tab. 3, gene lists in Suppl. Tab. 5): 1) Metabolic/housekeeping/proliferative (modules 2, 8, 9, 10, 13); 2) ECM/Migratory/Cytoskeletal (modules 1, 5, 11, 12); 3) Lineage and patterning (modules 6 and 7); 4) Axial mesoderm (module 3). Aggregate expression of the top 20-scoring hub genes for each of the modules showed altered expression in regulatory mutants compared to WT cells (Tab. 3) but, given the known *in vivo* roles of *Tbx1* in development and disease (Baldini et al. 2017), we focused our attention on the modules that were characterized by expression of genes involved in lineage determination and patterning (modules 6 and 7). Module 6 was characterized by genes involved in CPM lineage specification and includes the *Tbx1* gene among the top scoring hub genes. Aggregate expression of the 20 top-scoring hub genes of module 6 was significantly down regulated in ΔPRS10 and ΔPRS12 cells (Table 3 and Fig. 7b) compared to WT cells, while we found no significant differences between ΔPRS11 and WT cells. Reduced expression of a cardiopharyngeal gene network in cells with significant reduction of *Tbx1* gene expression is consistent with our previous findings (Lanzetta et al. 2025) and suggests that the deletion of PRS10 or PRS12 is akin to *Tbx1* partial loss of function. We validated the down regulation of module 6 against a published dataset of mouse embryonic tissue scRNA-seq (*Mesp1^Cre^*-driven deletion of *Tbx1* in E9.5 embryos (Nomaru et al. 2021)), (*P*=5.1×10^-6^, Wilcoxon rank sum test with continuity correction). Less predictable was the finding of significant up regulation of module 7 in all deletion mutants. This module is characterized by hub genes involved in anterior-posterior patterning and presomitic mesoderm specification, including genes involved in retinoic acid signaling. We were surprised by the up regulation of genes that are normally expressed in the paraxial/presomitic/somitic mesoderm, which is mainly posterior to the *Tbx1* gene expression domain, typically in the branchiomeric/anterior mesoderm. Also in this case, we confirmed the up regulation using the mouse tissue datasets mentioned above (*P*=0.002). A top scoring gene in module 7 is *Meox1*, which encodes a transcription factor known for its role in regionalization of paraxial/somitic mesoderm and differentiation (Candia et al. 1992; Mankoo et al. 2003; Skuntz et al. 2009). *Meox1* was mainly expressed in cluster S-0 of ΔPRS10 and ΔPRS12 cells, a cluster that is relatively less well differentiated according to RNA-velocity assay, and was barely detectable in WT cells (Fig. 8a). There was little or no overlap with *Tbx1* gene expression and in the deleted mutants it was strongly expanded and mostly located in cells of cluster S-0 (avg. log2FC=2.8, adj *P*<<10^−6^ in ΔPRS10 cells; avg. log2FC=3.1, adj *P*<<10^−6^ in ΔPRS12 cells). A similar observation concerned the gene *Hoxb3*, which is also involved in somite development (Manley and Capecchi 1997). The UMAP pattern of *Hoxb3* expression is similar to that of *Meox1* (Fig. 8b). Intrigued by these observations, we performed *in situ* hybridization of *Meox1* and *Hoxb3* on mouse embryos. In WT E9.5 embryos, *Meox1* was strongly expressed in somites, and in the mesenchyme of the 2^nd^ and 3^rd^ pharyngeal arch (PA) (Fig. 9a-d). Partial expression overlap with *Tbx1* was limited to the mesenchyme of the 3rd PA. In *Tbx1* dosage mutant embryos *Tbx1^neo2^*^/neo2^ and *Tbx1^neo2^*^/-^, which express 40% and 20% *Tbx1* RNA, respectively, we found expression in the head mesenchyme, particularly in the posterior periotic mesenchyme, a region where *Meox1* is normally expressed at a very low level, if at all (Fig. 9a’, b’, c’). In *Tbx1^neo2^*^/-^ (Fig. 9a’’, b’’, c’’) and *Tbx1*^-/-^ embryos (Fig. 9d’), *Meox1* was strongly expressed in the periotic region surrounding the entire otocyst. Additional images are shown in Suppl. Fig. 6. In contrast to E9.5 embryos, at E8.5 (Suppl. Fig. 7a) we found broad overlap between *Meox1* and *Tbx1* expression patterns in the head mesenchyme and the trunk (Suppl. Fig. 7b,c), a region similar to the one previously indicated as head-trunk interface (Dumas et al. 2024). In the *Tbx1*^-/-^ mutants we did not detect any changes in *Meox1* expression pattern (Suppl. Fig. 7a’, b’, c’). Thus, *Meox1* expression anomalies in *Tbx1* mutants arise between E8.5 and E9.5. A possible, speculative interpretation of this finding is that in E9.5 mutants there is persistence of the E8.5 pattern, with extensive expression in the head and trunk mesenchyme.

**Table 3.**
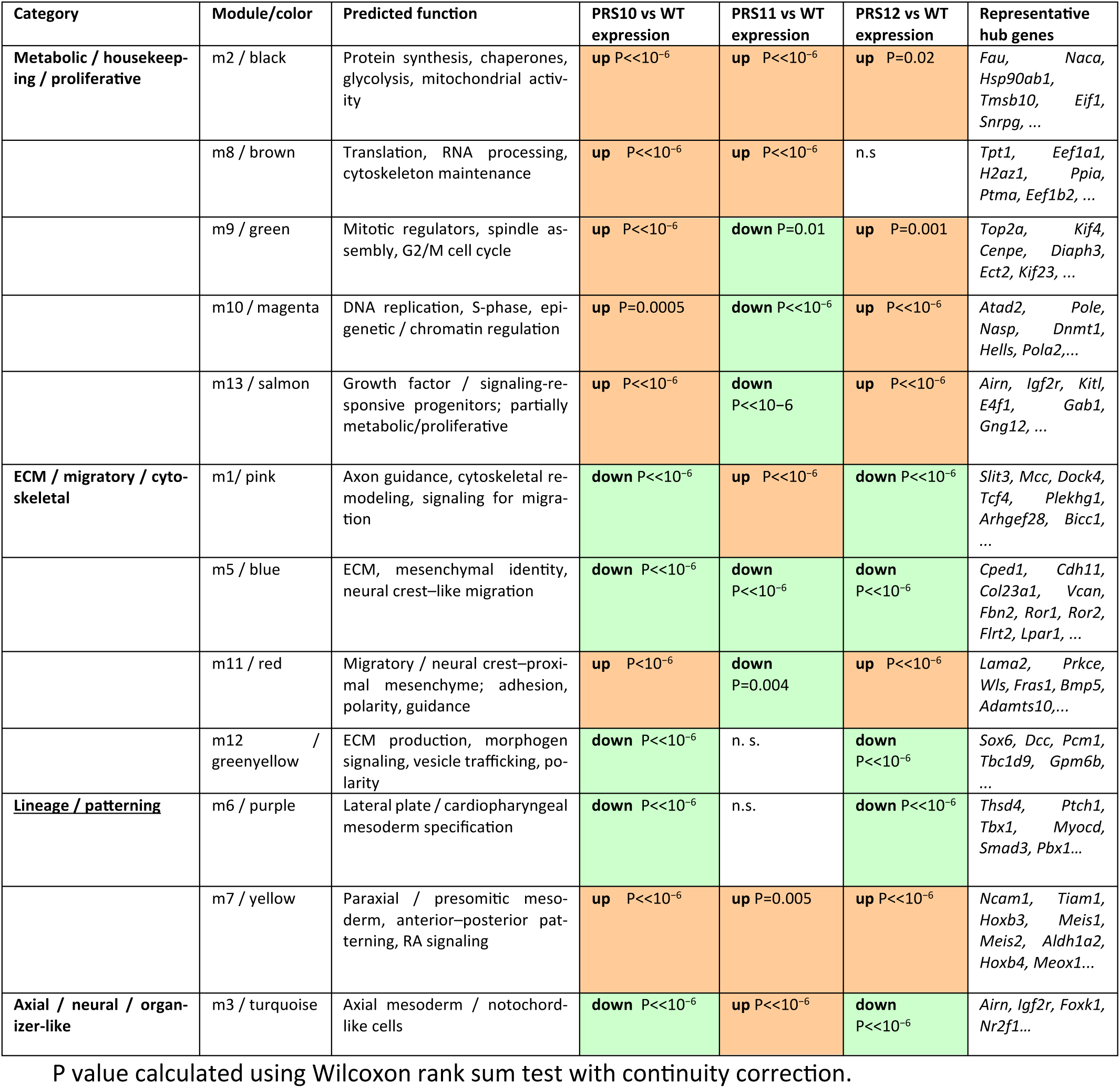
Transcriptional modules resulting from co-regulation analysis.

**Figure 7.**
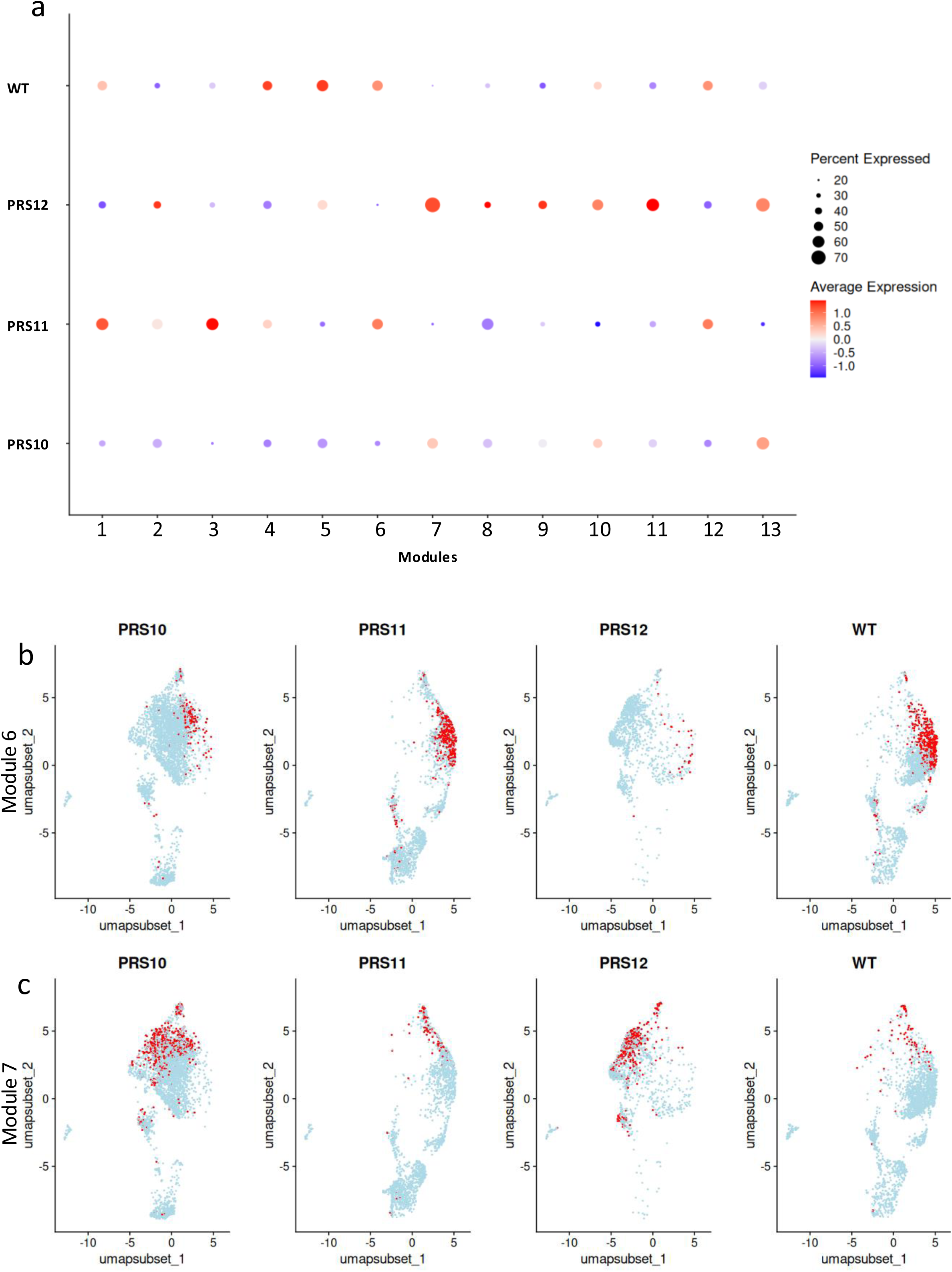
Co-regulation analyses identified transcriptional modules depleted or enriched in mutant cells. a) Dot plot illustrating the expression of transcriptional modules revealed by hdWGCNA analyses across genotypes. The color refers to the average level of expression while the diameter of the dots refers to the percentage of cells expressing the modules. The definition of the modules is listed in Table 3; b) aggregate expression of module 6 genes, which include *Tbx1*, predicted to be involved in cardiopharyngeal mesoderm lineage specification; c) aggregate expression of module 7 genes predicted to be involved in anterior-posterior patterning and presomitic mesoderm specification.

**Figure 8.**
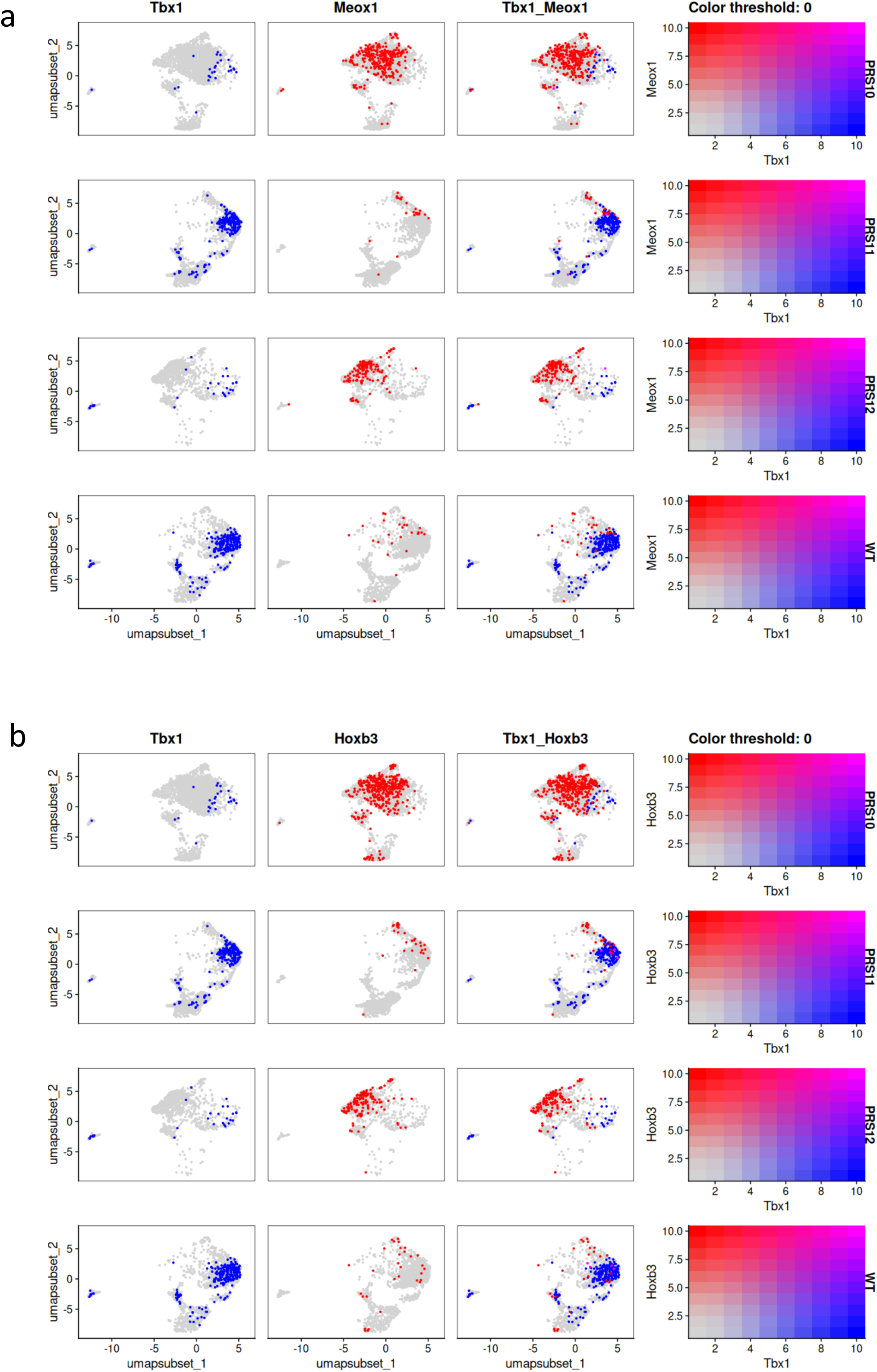
Module 7 genes Meox1 and Hoxb3 are mostly expressed in mutants ΔPRS10 and ΔPRS12. a) Dual-color UMAP display of *Meox1* and *Tbx1* gene expression. The expression pattern of the two genes is mostly non-overlapping in this differentiation model. *Meox1* expression is most evident in cluster S-0, the least differentiated cell population; b) *Hoxb3* gene expression exhibits similar characteristics as *Meox1* expression.

**Figure 9.**
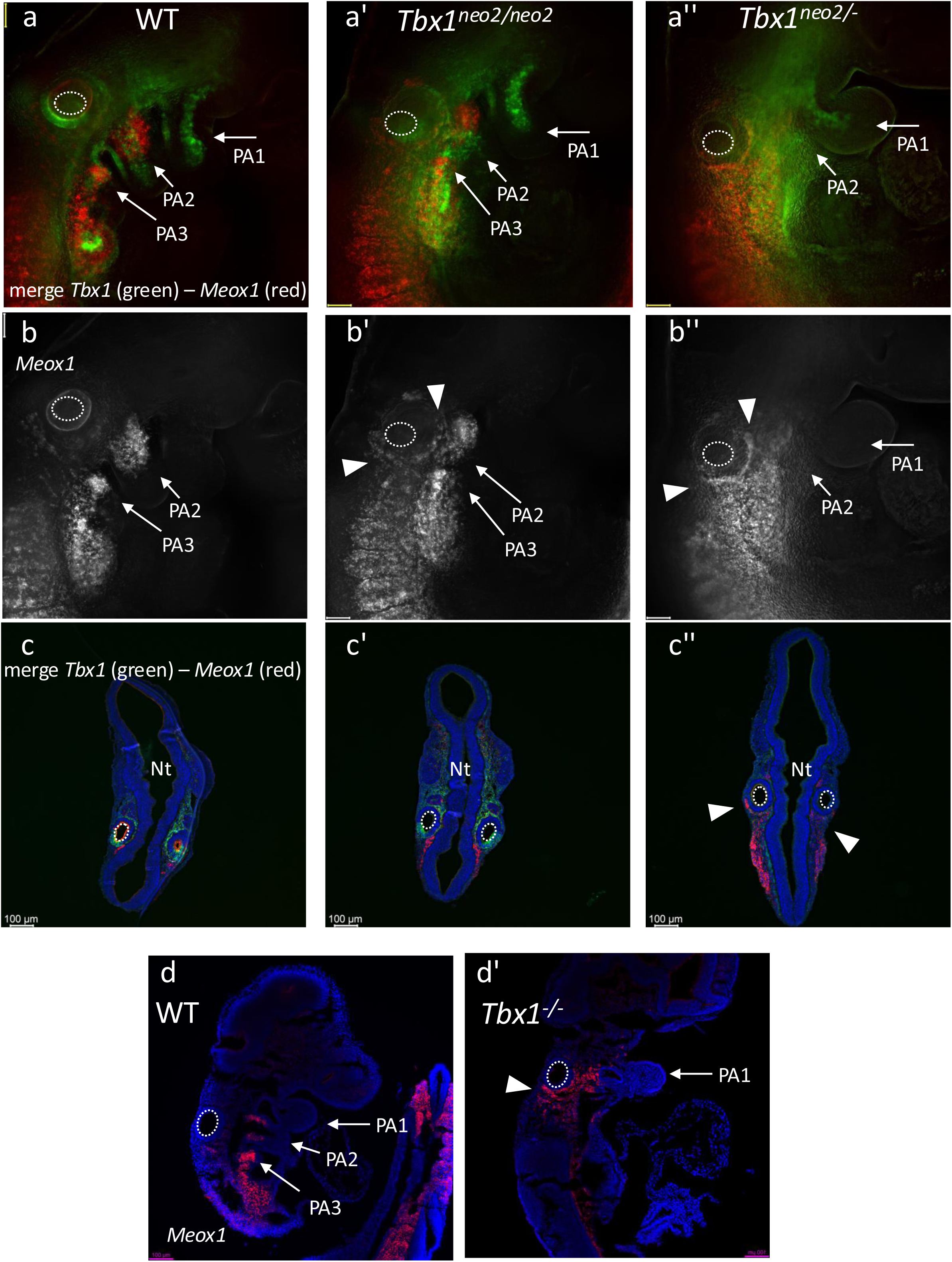
Meox1 gene expression is up regulated in the head/periotic mesenchyme of E9.5 mouse embryos carrying Tbx1 hypomorphic and null mutations. Dual color *in situ* hybridization (RNAscope) experiments using *Meox1* (red) and *Tbx1* (green) probes. a, a’, a’’) Images of whole mount embryos (right side), WT, *Tbx1^neo2/neo2^* (expressing approx. 40% of wild-type mRNA level), and *Tbx1^neo2^*^/-^ (expressing approx. 20% of wild-type mRNA level) embryos; b, b’, and b’’ are the same images but show only the *Meox1* signal; c, c’, c’’ show coronal sections of the same embryos at the level of the otocyst; d) sagittal sections of a E9.5 WT embryo hybridized with a *Meox1* probe compared to a similar section of a *Tbx1*^-/-^ embryo at the same stage (d’). Arrowheads indicate the ectopic *Meox1* signal in mutants. In all images the internal lumen of the otocyst is indicated for reference as dashed circles. PA1, 2, 3: arrows indicate the position of pharyngeal arch 1, 2, 3, respectively. Nt: neural tube. Scale bar: 100µM.

*Hoxb3* expression was mainly detected in the peripharyngeal mesenchyme/paraxial mesoderm of the posterior pharynx and somites and in the third and fourth PAs of WT embryos at E9.5 (Supplementary Fig. 8a-c). In *Tbx1*^-/-^ E9.5 embryos, the peripharyngeal domain is essentially undetectable with, instead, a broad expression in a more anterior region closer and partially surrounding the otocyst (Suppl. Fig. 8a’-c’ and Suppl. Fig. 9a’, b’, c’).

### Inference of a Regulatory hierarchy

Using a machine-learning tool implemented in the hdWGCNA package (Childs et al. 2024; Morabito et al. 2023) (function: ConstructTFNetwork) we crossed WT transcriptomic co-regulatory data with transcription factor binding sites in order to infer the transcription factor->targets pairings that might be at play in our cell differentiation model. The inferred top scoring regulators of *Tbx1* are shown on Tab. 4. By far the most high scoring candidate was a EBF transcription factor, a known, evolutionary conserved contributor to cardiopharyngeal specification (Razy-Krajka et al. 2014) and a marker of CPM (Argiro et al. 2024). Previously, we found *Ebf* genes in a transcriptional module affected by *Tbx1* mutation (module c3mod4 down regulated in the mutant and linked to somatic mesoderm and anterior cardiopharyngeal mesoderm) (Lanzetta et al. 2025). Also, among the top 5 potential regulators was FOXC2, previously reported as a *Tbx1* gene regulator (Maeda et al. 2006; Yamagishi et al. 2003). Using a bioinformatic tool (FIMO v. 5.5.9, 2025) (Grant et al. 2011) we scanned the PRS10, 11, and 12 sequences for the occurrence of high-confidence transcription factor binding sites (Supp. Tab. 6). Intersecting these putative binding motifs with those of predicted transcription factors, we found hits for four out of five predicted regulators of the *Tbx1* gene (Tab. 4). As for downstream potential targets of TBX1, we intersected the DEGs detected in ΔPRS10 and ΔPRS12 cells (n=607 unique genes) with genes annotated with TBX1 ChIP-seq peaks obtained using mouse embryo tissue (Nomaru et al. 2021) (n=444 genes annotated to 255 peaks) and obtained 44 genes that are listed in Suppl. Table 7. The list includes genes involved in several developmental pathways previously known to be related to the developmental functions of *Tbx1*, for example RA signaling genes, *Nrg1*, *Six1*, *Tshz2, Fn1, Aldh1a2, Ror1* and others (Suppl. Table 7).

**Table 4.** Inferred regulators of *Tbx1* gene transcription.

|  | tf | gene | Gain | Cover | Frequency | Cor | FIMO* Seq<br>match | pVal<br>(FIMO*) | Occurrences |
| --- | --- | --- | --- | --- | --- | --- | --- | --- | --- |
| 1 | Ebf2 | *Tbx1* | 0.542 | 0.168 | 0.168 | 0.616 | PRS10, 11,<br>12 (EBF) | <1e-04 | 17 (PRS10<br>n=12; PRS11<br>n=1; PRS12<br>n=4) |
| 2 | Glis1 | *Tbx1* | 0.170 | 0.128 | 0.128 | 0.511 | none | n.a. | 0 |
| 3 | Zbtb14 | *Tbx1* | 0.050 | 0.068 | 0.068 | 0.319 | PRS11 | 3.3e-06 | 1 |
| 4 | Fosl2 | *Tbx1* | 0.022 | 0.034 | 0.034 | 0.155 | PRS10<br>(FOS:JUN) | 6.5e-05 | 1 |
| 5 | Foxc2 | *Tbx1* | 0.021 | 0.054 | 0.054 | 0.150 | PRS10, 11,<br>12 (FOX) | <1e-04 | 29 (PRS10<br>n=10; PRS11<br>n=5; PRS12<br>n=14) |

*Shaded:* \*FIMO v. 5.5.9, 2025 scan.

Thus, in the differentiation model used here, reduced dosage of *Tbx1* expression might be driven by EBF transcription factors, which are already known for their role in cardiopharyngeal mesoderm (Razy-Krajka et al. 2014). Downstream, we observed reduced expression of a cardiopharyngeal specification transcriptional module and up regulation of a transcriptional module involved in embryonic patterning. The cartoons shown in Fig. 10 summarize the main results of this work.

**Figure 10.**
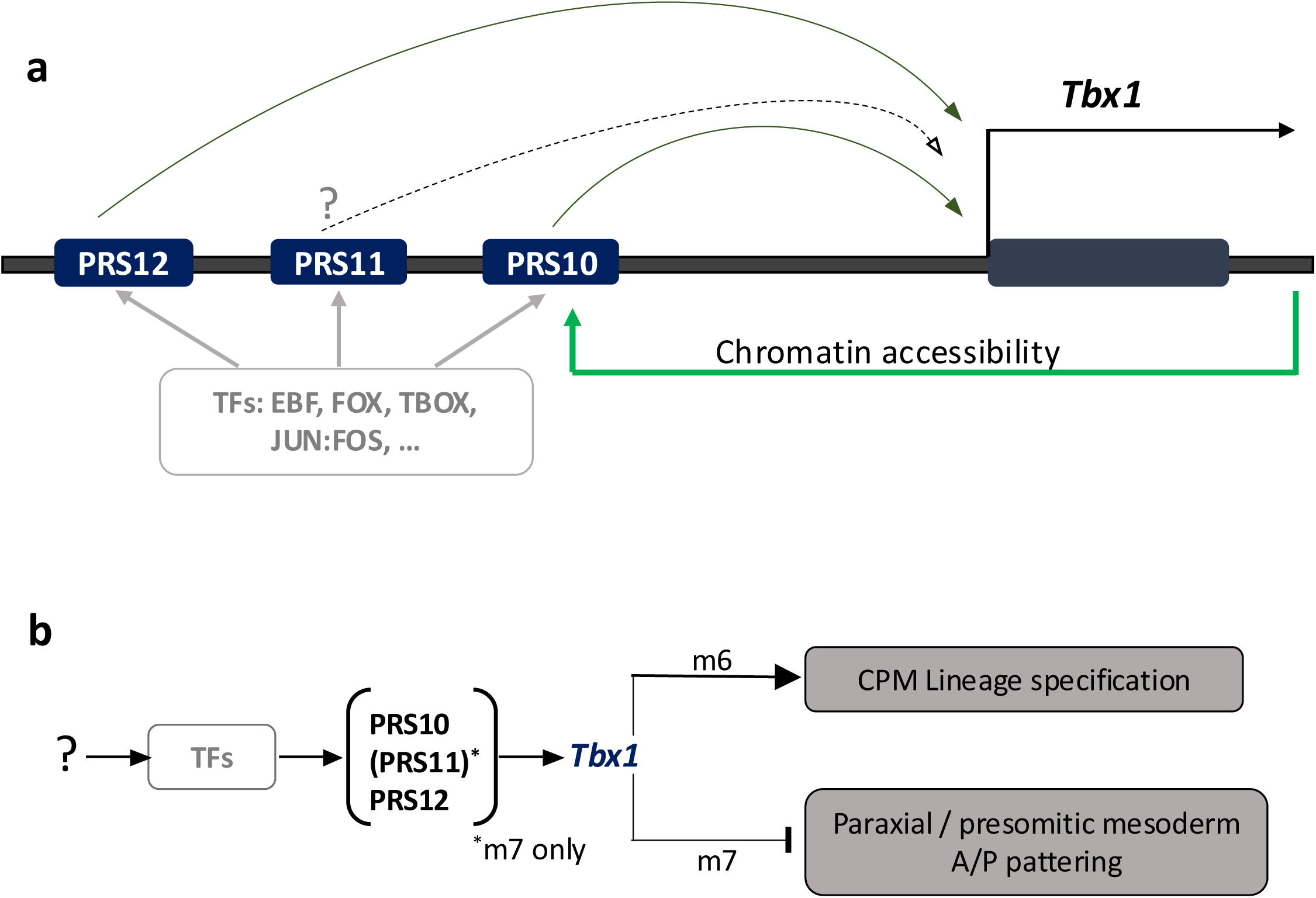
Schematic summary. a) Cartoon of the interactions between transcription factors (TFs, gray) and the cluster of putative regulatory sequences (PRSs) located upstream of the *Tbx1* gene. The dotted line indicates uncertainty over the role of PRS11. The green line indicates a feedback loop affecting chromatin accessibility of the PRS10 segment. b) PRS10 and PRS12 drive *Tbx1* gene expression to regulate positively a CPM lineage specification pathway, while also driving the suppression (along with PRS11) of anterior/posterior (A/P) patterning genes. The question mark refers to unknown signals or transcription factors upstream to the *Tbx1* regulators.

## DISCUSSION

In this report we implemented a strategy to identify regulatory sequences starting from chromatin accessibility data generated with the transposase-based technique ATAC-seq (Buenrostro et al. 2015) combined with simultaneous transcriptional information from single nuclei. Training a computer to score potential enhancers based on epigenomic data deposited in public databases offered a relatively simple approach to score ATAC-seq peaks for their probability to be enhancers. This approach led to the identification of a cluster of putative regulatory sequences at the *Tbx1* locus that were then validated by gene editing and gene expression analyses, including single-cell assays.

Comparison of the genomic coordinates of the cluster with literature data revealed that PRS10 overlapped partially with a previously identified FOX-containing regulatory sequence (Brown et al. 2004; Yamagishi et al. 2003) and it was also included in a genomic segment that we have previously deleted in mice (Zhang and Baldini 2010). The former studies were based on classical transgenic analyses with a reporter vector, while the latter was based on endogenous sequence deletion, thus more directly comparable to the study presented here. In the mouse embryo, the deletion of this genomic segment led to an approx. 25% reduction of *Tbx1* gene expression (using quantitative whole-embryo assays), but no morphological phenotypic anomalies (Zhang and Baldini 2010). This is in apparent contrast with the much deeper suppression of gene expression associated with PRS10 deletion in our differentiation model. Differences may be due to a more efficient compensation mechanism during embryonic development than in the cellular model, perhaps mediated cell non-autonomously by tissues or cell types that are not present in the cell culture system. We found that the deletion of PRS10 or PRS12 did not structurally prevent transcription of the gene because we detected subsets of cells that expressed *Tbx1* in both deletion mutants.

One reason for performing this study was to address mechanisms of *Tbx1* haploinsufficiency. It is reasonable to speculate that mRNA dosage reduction will lower the speed at which the protein accumulates and thereby impair timely adaptation to molecular cues in morphogenetic and differentiation processes. Thus, from this perspective, haploinsufficiency is rooted in transcriptional regulation mechanisms (Veitia 2002). More specifically, a theoretical model (Zug 2022) predicted that a gene encoding a transcription factor is likely to be haploinsufficient if it possesses three key features: i) has a function in cell fate determination, ii) is part of a positive feedback regulatory loop, and iii) is controlled by a super-enhancer-like cluster of regulatory sequences. A role for *Tbx1* in governing differentiation trajectories has been shown in various models (Dastjerdi et al. 2007; Kong et al. 2014; Lanzetta et al. 2025; Nomaru et al. 2021; Wang et al. 2019), including in the study presented here. The cluster CLRE identified here may be considered a super-enhancer-like region as it includes at least three regulatory sequences (ATAC peaks are relatively large and may include multiple enhancers) within a relatively small region, thus it may be seen as a hub for interaction of many transcription factors. In addition, we showed here that CLRE also included an element of positive feedback loop revealed by reduced chromatin accessibility in the absence of *Tbx1* gene expression. Therefore, the data presented here, and published data are compatible with a model in which the architecture of *Tbx1* gene regulation predisposes to haploinsufficiency. While additional data will be required to prove this hypothesis, we think that it is important to consider it as it may provide a target to neutralize this important disease mechanism in the context of the 22q11.2 deletion syndrome.

In summary, our study identified a cluster of regulatory sequences required for the expression of *Tbx1* in a mesodermally-orientated cell differentiation model. Two of these sequences (PRS10 and PRS12) are required for the level of gene expression across the spectrum of cell types obtained with this model, although a small portion of mesodermal cells appear to be non-responsive. The transcriptomic consequences of individual deletions of these two regions are interpretable as a partial loss of function as we found impairment of a genetic network related to CPM specification, which is also affected by loss of function of *Tbx1*. The up regulation of anterior/posterior patterning genes such as *Meox1* and *Hoxb3,* which was observed when all three PRSs were individually deleted and in *Tbx1* mutant mouse embryos, was a surprise finding that warrants further investigation. Here, we speculate that there may be a cell intrinsic mechanism that underlies the very severe defects of the posterior pharyngeal apparatus that were identified in the first descriptions of *Tbx1* mouse mutants (Jerome and Papaioannou 2001; Lindsay et al. 2001; Merscher et al. 2001).

## MATERIALS AND METHODS

### Cell lines and differentiation model

ES-E14TG2a mouse embryonic stem cells (mESC) line (ATCC CRL-1821) were cultured without feeders and maintained undifferentiated on gelatine-coated dishes in GMEM (Sigma Cat# G5154) supplemented with 103 U/ml ESGRO LIF (Millipore, Cat# ESG1107), 15% fetal bovine serum (ES Screened Fetal Bovine Serum, US Euroclone Cat# CHA30070L), 0.1 mM non-essential amino acids (Gibco, Cat# 11140-035), 0.1 mM 2-mercaptoethanol (Gibco, Cat# 31350-010), 0.1 mM L-glutamine (Gibco, Cat# 25030081), 0.1 mM Penicillin/Streptomycin (Gibco, Cat# 10378016), and 0.1 mM sodium pyruvate (Gibco, Cat# 11360-070). The cells were passaged every 2–3 days using 0.25% Trypsin-EDTA (1X) (Gibco, Cat# 25200056) as the dissociation buffer.

For precardiac organoids (POs) differentiation, wild type and mutant cells were dissociated with Trypsin-EDTA and plated at a density of 100,000 cells/ml in serum-free differentiation media made of 75% Iscove’s modified Dulbecco’s media (Cellgro, 15-016-CV) and 25% HAM F12 media (Cellgro, 10-080-CV). The media were supplemented with N2 (Gibco, 17502048) and B27 (Gibco, 12587010), penicillin/streptomycin (Gibco, 10378016), 0.05% bovine serum albumin (Invitrogen, P2489), L-glutamine (Gibco, 25030081), 5 mg/ml ascorbic acid (Sigma-Aldrich, A4544) and 4.5×10−4 M monothioglycerol (Sigma-Aldrich, M-6145). After 48 h, the POs were harvested and seeded in serum-free differentiation media supplemented with 1.5 ng/ml human activin A (Peprotech, 120-14E) and 1.5 ng/ml human BMP4 (Peprotech, 120-05ET). Media were subsequently changed to serum-free differentiation media without additional growth factors after 2 days, and the POs cultured until d6 or d8. EBs were left to settle to the bottom of 15 ml tubes (∼2-5 min) before removing the media and the POs were resuspended in DPBS without calcium and magnesium. POs were disaggregated using the StemPro® Accutase® disaggregation kit (Thermo Fisher Scientific, A1110501) for 4-8 min at room temperature, to snRNAseq, and using Embryoid Body Dissociation Kit (Miltenyi Biotec, 130-096-348) for 10 min at room temperature to FACS analysis. The disaggregation process was blocked by adding fresh pre-warmed complete medium. Cells were then washed multiple times in DPBS without calcium and magnesium, and processed as follows. Total RNA was isolated from wild type and mutant POs derived cells with QIAzol lysis reagent (Qiagen #79306), according to the manufacturer’s protocol. The isolated RNAs were quantified using a NanoDrop spectrophotometer 1000. Before reverse transcription, RNA samples were treated with DNAse I to eliminate any contamination with genomic DNA (located in the interphase during extraction). cDNA was transcribed using 1 or 2 μg total RNA with the High-Capacity cDNA reverse transcription kit (Applied Biosystem catalog. n. 4368814). Quantitative gene expression analyses (qRT-PCR) were performed using SYBR Green PCR master mix (Applied Biosystem) with StepOnePlus™ Real-Time PCR System. The run used was similar to PCR default condition, but the number of cycles is increased up to 40 cycles. The cycle threshold (Ct) was determined during geometric phase of the PCR amplification plots, as illustrated in the manufacturer’s protocol. Primers are listed in Supplementary Table 1. Relative gene expression was evaluated using the 2-ΔCt method, and *Gapdh* expression as normalizer. Graph Pad Prism software v8.00 (GraphPad) was used to analysis of data. Relative mRNA levels were analyzed in triplicate, and data were presented as means ± SEM and ± SD. Two-way and one-way ANOVA with Tukey’s multiple comparisons test were used to assess that there is a statistically significant interaction effect between the two independent groups, “wild type” and “mutants” on a gene expression variable (Brown-Forsythe and Bartlett’s test). Differences were considered significant at p-value <0.05.

### Gene editing

CLRE and individually PRS deletions were induced in E14-Tg2a using Alt-R™ CRISPR-Cas9 System (IDT) following the manufacturer’s specifications. This genome editing system is based on the use of a ribonucleoprotein (RNP) consisting of Alt-R S.p. Cas9 nuclease complexed with an Alt-R CRISPR-Cas9 guide RNA (crRNA:tracrRNA duplex). The crRNA is a custom synthesized sequence that is specific for the target and contains a 20 nt sequence that is complementary to the tracrRNA. Alt-R CRISPR-Cas9 tracrRNA-ATTO 550 (5 nmol catalog no. 1075927) is a conserved 67 nt RNA sequence that is required for complexing to the crRNA to form the guide RNA that is recognized by S.p. Cas9 (Alt-R S.p. Cas9 Nuclease 3NLS, 100 μg catalogue no. 1081058). The fluorescently labeled tracrRNA with ATTO™ 550 fluorescent dye is used to FACS-purify transfected cells. The protocol involves three steps: (1) annealing of the crRNA and tracrRNA, (2) assembly of the Cas9 protein with the annealed crRNA and tracrRNAs, and (3) delivery of the ribonucleoprotein (RNP) complex into mESC by reverse transfection. Briefly, we annealed equimolar amounts of resuspended crRNA and tracrRNA to a final concentration (duplex) of 1 μM by heating at 95°C for 5 min and then cooling to room temperature. The RNA duplexes were then complexed with Alt-R S.p. Cas9 enzyme in OptiMEM media to form the RNP complex, which was then transfected into mESCs using the RNAiMAX transfection reagent (Invitrogen). After 48 h incubation cells were purified by Fluorescence Activated Cell Sorting (FACS). In brief, for gRNA transfection efficiency assay to detect and visualize the fluorescently labeled gRNA complex (crRNA:tracrRNA-ATTO™550), we dissociated mESC with Trypsin-EDTA and analyzed the % of fluorescence using the BD FACS ARIAIII™ cell sorter. Cells containing the transfected gRNA complex were individually isolated and plated in 96 multi-wells to obtain colony growth. We screened by PCR around 200 clones for each deletion: gDNAs were amplified using myTaq™ DNA polymerase (Meridian Bioscience) and a standard 3-step cycling PCR profile (10 min at 94°C, 30 amplification cycles with denaturation at 94°C for 30sec, annealing at 60°C for 30sec, and extension at 72°C for 30sec), followed by a final extension at 72°C for 10 min. The amplified products were separated on agarose gels and visualized by ethidium bromide staining. Positive clones were confirmed by DNA Sanger sequencing (Eurofins Genomics). Sequences of the gRNAs used and the PCR primers for each deletion are listed in Supplementary Table 1.

### Enhancer search and scoring

The RNA-seq and ATAC-set datasets integration and the subsequent enhancers prediction were performed as described (Aurigemma et al. 2024; Lanzetta et al. 2025). In brief, chromatin accessibility data from snATAC-seq was correlated with gene expression of *Tbx1* from snRNA-seq in the differentiated cell populations that constitute the precardiac organoids (datasets downloaded from the European Nucleotide Archive under accession number PRJEB64050 and analysed as described in Lanzetta et al., 2025). We have identified 14 ATAC peaks in 40 Kb genomic region of *Tbx1* locus. Sequences of these peaks were screened using a machine-learning method, trained on validated enhancers. We downloaded the coordinates of validated enhancers (i.e., positive regions with response Y=1) from the VISTA ENHANCER Browser. We randomly generated non-enhancers’ coordinates with the same cardinality and size (i.e., negative regions with response Y=0). We used available epigenetic data to build the feature matrix X by assessing the overlaps with the positive and negative regions. We trained the regression model using k-fold cross-validation. Then, we classified a region as a potential enhancer using the threshold of 0.5 over the average estimation of the expected probability of being an enhancer.

We used a publicly available dataset (GEO database number GSE198561) (Ranade et al. 2022) to test coverage of ATAC peaks of our PRS sequences in mouse embryo tissues. The dataset includes snATAC-seq from WT and *Tbx1*^-/-^ mouse embryos at E9.25. We downloaded samples and called peaks for each sample independently using MACS2 (v.2.2.9.1) and retained only standard chromosomes after removing ENCODE mm10 blacklist regions obtained from AnnotationHub (AH104614; snapshot date: 2025-04-08). We generated a consensus peak set across all samples and filtered peaks by length (20–10,000 bp). We quantified fragment counts per cell using a feature matrix built from each sample’s fragment file, retaining cells with at least 1,000 total counts, TSS enrichment >2, and nucleosome signal <4. We merged samples into a combined ChromatinAssay object and performed dimensionality reduction via TF-IDF normalization followed by singular value decomposition (LSI). To correct for batch effects across replicates, we integrated LSI embed-dings using anchor-based integration (FindIntegrationAnchors, IntegrateEmbeddings) and computed a UMAP from the integrated reduction. We identified clusters using a shared nearest-neighbour graph (Louvain algorithm, resolution 0.2).

### snRNA-seq

Wild type and mutant organoids on day 6 of differentiation were collected and disaggregated using the StemPro® Accutase® disaggregation kit (Thermo Fisher Scientific, A1110501) for 4-8 min at room temperature. The disaggregation process was blocked by adding fresh pre-warmed complete medium. Cells were then washed multiple times in DPBS without calcium and magnesium, and processed following 10X Genomics Chromium protocols (CG000124, CG000169). In brief, we treated the samples with lysis buffer and washing solutions to isolate nuclei from the other cellular components. We first assessed the successful outcome of cell lysis by microscope observation after staining with Trypan blue (around 90% of total). Then, isolated nuclei were incubated with RFP fluorescent dye and subjected to automated counting using Countess II FL Automated Cell Counter (ThermoFisher), to distinguish intact nuclei from damaged ones. We captured 10.000 nuclei from each sample, in two replicates, and loaded into the Chromium Controller (10X Genomics), to generate Gel Beads in emulsion for single nuclei. The library preparation for sequencing was carried out with the Chromium Single Cell 3’ reagents according to the manufacturer protocol. DNA purification and the size selection steps were done with SPRIselect magnetic beads (Beckman Coulter, Brea, CA) as indicated by the 10X protocol. Qubit dsDNA BR Assay (Thermo Fisher Scientific, USA) were used to determine the concentration of each indexed library, and quality control were assessed using D1000 screentape on the TapeStation 4150 (Agilent Technologies, Santa Clara, CA). DNA libraries were sequenced into the Illumina NextSeq550 System (Illumina, San Diego, CA) in a paired-end dual index format with the following sequencing cycles: 28 for Read1, 10 for Index i7, 10 for Index i5 and 90 for Read2. We processed Fastq files from each 10x Genomics run using the Cell Ranger (v. 8.0.1) pipeline for demultiplexing and gene alignment. We imported the resulting count matrices into R (v. 4.5.3) and analysed them using the Seurat package (v. 5.4.0). We first performed quality control independently for each sample. We retained cells with more than 500 and fewer than 20,000 total counts, between 500 and 3,000 detected genes, less than 5% mitochondrial gene content, and less than 30% ribosomal gene content. We removed mitochondrial genes, ribosomal genes, and *Malat1* from the expression matrix prior to downstream analyses. We then applied the standard Seurat workflow to each sample. We log-normalized the raw counts (scale factor = 10,000), identified the top 3,000 highly variable genes using the “vst” method, scaled the data, and performed dimensionality reduction using principal component analysis (PCA) with 30 components. We constructed a shared nearest neighbor graph and performed clustering at multiple resolutions.

We identified and removed potential doublets using DoubletFinder (v.2.0.6). The optimal pK parameter was determined through a parameter sweep and maximization of the BCmetric. The expected number of doublets was estimated assuming a 7.5% doublet formation rate and adjusted for homotypic doublets. We excluded doublets from further analyses.

We then merged all samples into a single Seurat object. The merged dataset was log-normalized, and 2000 highly variable genes were selected. We scaled the data while regressing out total UMI counts (nCount_RNA) and performed PCA. We corrected the batch effects across samples using the Harmony integration method implemented in Seurat. We then performed dimensionality reduction using the Harmony embeddings and retained the first 30 Harmony components for downstream clustering and visualization. We chose clustering performed at resolution 0.3, and UMAP was computed in the integrated space. We identified cluster-specific marker genes using the FindAllMarkers function (logfc.threshold = 0.25, min.pct = 0.25) and used the top-ranking markers for cluster annotation. We quantified gene signature activity at the single-cell level using the AddModuleScore function Seurat v5. For each predefined gene set, AddModuleScore computed a per-cell module score by averaging the normalized expression values of the genes in the signature and subtracting the aggregated expression of randomly selected control gene sets matched for expression bin distribution. The resulting module scores were used to evaluate relative enrichment of specific transcriptional programs across cellular subpopulations and experimental conditions.

We selected clusters 2, 9, and 10 (defined at resolution 0.3 in the Harmony-integrated space) for downstream subclustering analysis. We subsetted these clusters and reprocessed them independently. For this analysis, we performed normalization, identification of 2000 highly variable genes, scaling, and PCA. We retained the first 20 principal components for downstream dimensionality reduction, neighbourhood sgraph construction, clustering, and UMAP visualization. We conducted differential expression comparisons on the subset independently between the PRS10, PRS11, and PRS12 conditions against the WT reference group using the FindMarkers function. We performed Gene-level statistical testing using the Wilcoxon rank-sum test (test.use = “wilcox”). We considered only genes expressed in at least 10% of cells in either comparison group (min.pct = 0.1) and exhibiting a minimum log fold-change threshold of 0.2 (logfc.threshold = 0.2). We filtered resulting marker tables to retain significantly differentially expressed genes with adjusted p-values (p_val_adj) below 0.01 after multiple testing correction.

### Multidimensional Scaling

To assess the Euclidean distance between the transcriptomes of the different genotypes in a reduced-dimensional space, we applied a pseudobulk approach and generated a multidimensional scaling (MDS) plot using muscat (v1.14.0) (Crowell et al. 2020). An MDS plot was then computed, where each centroid represents each replicate of each genotype (WT, ΔPRS10, ΔPRS11, ΔPRS12).

### Network analysis

To identify transcriptional modules associated with the WT genotype, we performed a high-dimensional Weighted Gene Co-expression Network Analysis using the hdWGCNA R package (0.4.08) on the Seurat subset. To mitigate technical noise and dropout effects, we performed metacell aggregation using the MetacellsByGroups function. We identified k=25 nearest neighbours in the Harmony latent space, allowing a maximum of 15 shared cells between metacells (max_shared = 15). For network construction, we retained genes expressed in at least 5% (fraction = 0.05) of the cells within the analysed subset.

We constructed the co-expression network for the WT group using a signed adjacency matrix. We selected a soft-thresholding power based on scale-free topology criteria. We identified the modules via hierarchical clustering using the following parameters: minModuleSize = 15, deepSplit = 3, and mergeCutHeight = 0.25. we calculated Module Eigengenes (MEs) and connectivity (kME) to identify intramodular hub genes.

### Motif Scanning and Transcription Factor Network Inference

To identify regulatory drivers, we performed a motif scanning analysis using the JASPAR2024 database (vertebrate CORE collection) (v. 0.99.7) and the motifmatchr package (v. 1.30.0). Position Frequency Matrices (PFMs) were mapped to the mm10 mouse genome (via EnsDb.Mmusculus.v79 (v.2.99.0). We standardized the transcription factor (TF) names to match the Seurat object nomenclature. We inferred a TF-regulatory network using an XGBoost regression model (extreme gradient boosting). We configured the model with the following parameters: objective = “reg:squarederror”, max_depth = 1, eta = 0.1, and alpha = 0.5. This supervised learning approach was used to predict target gene expression based on TF expression levels.

### Regulon Discovery and Enrichment

We identified and categorized TF-target interactions (regulons) using two distinct strategies: A) Prioritizing TFs by selecting the top 10 target genes per TF B) Prioritizing genes by selecting the top 50 TFs per target gene. We applied a regulatory threshold of 0.01 to both strategies to filter significant interactions. Finally, we exported the inferred TF-target connections for downstream validation against differentially expressed genes (DEGs).

To identify and score putative TF binding sites within the PRS10, PRS11, and PRS12 sequences, we used the FIMO tool online (https://meme-suite.org/meme/doc/fimo.html v. 5.5.9) (Grant et al. 2011) with default parameters and the JASPAR2022_CORE_non-redundant_pfms_meme database.

### RNA velocity analysis

We performed RNA velocity analysis in Python (v 3.10.19) using packages scanpy (v 1.11.5) and scVelo (v 0.3.3). on each genotype separately. Loom files containing spliced and unspliced transcript count matrices were imported into AnnData objects using anndata.read_loom. We standardized cell barcode identifiers by replacing terminal barcode suffixes and harmonizing sample-specific prefixes to ensure consistency across datasets.

We filtered the cells included in the analysis by matching the loom-derived cell barcodes with the metadata table containing the selected cell identifiers. We subsequently concatenated the filtered datasets into a single AnnData object for downstream analysis.

To preserve the original dimensional reduction generated during preprocessing, we imported the original umap coordinates calculated on Seurat. We also imported cluster annotations and added to the cell-level observations. We processed the combined dataset using the scVelo preprocessing workflow. Specifically, we filtered and normalized genes using scv.pp.filter_and_normalize with a minimum shared count threshold of 20 and we selected the top 3000 highly variable genes. We retained the gene *Tbx1* throughout the filtering process to ensure its inclusion in downstream analyses. We computed first- and second-order moments across neighboring cells using scv.pp.moments with 30 principal components and 30 nearest neighbours. We estimated RNA velocities using both the stochastic and dynamical models implemented in scVelo. Initially, we calculated velocities using the default stochastic framework with scv.tl.velocity, followed by construction of the velocity graph using scv.tl.velocity_graph.

To infer transcriptional dynamics at higher resolution, we applied the dynamical model implemented in scv.tl.recover_dynamics. We then recalculated RNA velocities using mode=’dynamical’ and reconstructed the velocity graph. We visualized velocity streams on the imported UMAP embedding using scv.pl.velocity_embedding_stream, with cells colored according to cluster identity.

### RNAscope

We collected mouse embryos collected at E8.5 and E9.5 (13 and 24-25 somites, respectively) obtained by crossing mouse lines *Tbx1^neo2^*^/+^ (Zhang et al. 2006) and *Tbx1^lacZ^*^/+^ (Lindsay et al. 2001) in a clean facility in a C57BL/6N background. All mouse experimentation was carried out according to a protocol approved by the Italian Ministry of Health (28/2022-PR to AB). Genotyping was carried out according to instructions provided by the original reports. Whole embryos were fixed in 4% PFA, dehydrated and rehydrated with increasing / decreasing concentrations of methanol and hybridizated with the mRNA probes according to the RNAscope™ ACD-Biotechne manufacturer’s instructions. Embryos were permeabilized for 20 min with Protease III (15 min for E8.5 embryos), followed by overnight incubation at 40°C with the probe solution. We used commercial probes provided by RNAscope: Probe-Mm-Meox1-C1 (530641); Probe-Mm-Tbx1-C2 (481911-C2); Probe-Mm-Hoxb3-C2 (515851-C2). The signal was developed with TSA-FITC (Akoya Bioscience, NEL741001KT; 1:500) and TSA-CY3 (Akoya Bioscience, NEL745001KT; 1:2000). Embryos were embedded in optimal cutting temperature (OCT) compound and cryosectioned sagittal and coronal at a thickness of 10 μm. Images were acquired with Leica Thunder Imaging System (Leica Microsystems).

## Supporting information

Supplementary fugures

## Acknowledgements

We thank Rosa Ferrentino and Giuseppina Divisato for technical support. We acknowledge the support of the Microscopy Core and the NGS core of the Department of Molecular Medicine and Medical Biotechnology, University of Naples Federico II, and the mouse facility of the Institute of Genetics and Biophysics of the CNR.

## Author contributions

SA: wet-bench experiments, conceptualization, manuscript editing; OL: Bioinformatic strategies and analyses; MB: mouse breeding and handling, immunofluorescence, RNAscope; PS: technical support with snRNA-seq; RF: mouse genotyping, cell culture. PZ: raw data handling, QC; GM: snRNA-seq supervision; CA: bioinformatic analysis supervision, manuscript editing; AB: project conceptualization, funding recruitment, manuscript writing.

## Data and resource availability

The snRNA-seq data are deposited to the GEO databank under accession number GSE334262. The mouse mutant lines used in this work are available through the EMMA/Infrafrontiers repository under codes EM:02136 (*Tbx1^neo2^*^/+^) and EM:02137 (*Tbx1^lacZ^*^/+^). All the other data are reported in the manuscript and the accompanying supplementary material.

## SUPPLEMENTARY INFORMATION

### TABLES

**Supplementary Table 1**

Primers and gRNA oligonucleotide sequences.

**Supplementary Table 2**

Top 20 markers of 11 clusters of the full dataset (from Fig. 4a).

**Supplementary Table 3**

A list of differentially expressed genes identified within clusters 2, 9, and 10 of the full dataset in comparisons mutant vs WT.

**Supplementary Table 4**

List of top 50 markers calculated within the subset re-clustering (clusters S-0 to S-5).

**Supplementary Table 5**

Top hub genes of transcriptional modules obtained using hdWGCNA co-regulation algorithm.

**Supplementary Table 6**

Predicted transcription factor binding sites of PRS10, 11, and 12 identified by FIMO scan.

**Supplementary Table 7**

Genes returned after intersection of differentially expressed genes (Supplementary Table 3) and genes annotated to the TBX1 ChIP-seq peaks of mouse embryos previously reported (Nomaru et al. 2021).

### FIGURES

**Supplementary Figure 1**

Coverage map of snATAC-seq data obtained from mouse embryos at E9.25 (Ranade et al. 2022) focused on our CLRE region. The PRS10, PRS11, and PRS12 peaks are easily identifiable. Note that some peaks (e.g. PRS10) appear split in some clusters, suggesting that they may include multiple enhancers.

**Supplementary Figure 2**

The CLRE is located within the same Topologically Associating Domain as the *Tbx1* promoter. HiC data obtained from data.4dnucleome.org.

**Supplementary Figure 3**

Reduced coverage of PRS10 in two clusters (c7 and c13) of a published dataset of snATAC-seq in *Tbx1*^-/-^ vs WT mouse embryos at E9.25 (Ranade et al. 2022).

**Supplementary Figure 4**

Strategy applied to gene editing of PRS sequences. a) Schematic representation of the CRISPR-Cas9 strategy b) differentiation protocol workflow. To achieve precise deletion of the three PRSs both in cluster and individually, we selected two crisprRNAs with binding specificity to direct the Cas9 endonuclease complex upstream and downstream of the targets; the ribonucleoprotein (RNP) complex was introduced into mESCs using transfection reagents (created using BioRender.com).

**Supplementary Figure 5**

a) deletion of the entire CLRE; b) deletion of PRS10; c) deletion of PRS11; and d) deletion of PRS12. In all cases we used two RNP complexes bound upstream and downstream of the sequence to be deleted. To test if the cleavages occurred, PCR reactions were performed with upstream and downstream primers designed to the target regions, as shown. For each panel we show FACS purification results of the pool of transfected cells (left), next there are examples of the PCR results obtained from pool of cells positive for transfection and on the right there are examples of PCR after of individual clones after isolation.

**Supplementary Figure 6**

Two-color RNAscope images of whole-mount E9.5 mouse embryos with the genotype indicated: *Meox1* is shown in red and *Tbx1* in green. In all panels, the dashed circle drawing indicates the lumen of the otocyst.

*Tbx1^neo2^*^/neo2^ embryos express approx. 40% of the WT *Tbx1* mRNA; *Tbx1^neo2/-^* (an abbreviation for *Tbx1^neo2^*^/lacZ^) express approx. 20% of the WT *Tbx1* mRNA. PA1, 2, 3: arrows indicate the position of pharyngeal arches 1, 2, or 3, respectively. Arrowheads indicate ectopic *Meox1* signal.

The scale bar is 100µm.

**Supplementary Figure 7**

Images of RNA-scope hybridization of whole mount E8.5 (13 somites) embryos (a and a’) using a *Meox1* probe, and transverse sections (b-c and b’-c’) of the same embryo showing also the *Tbx1* probe (in green). Note that the *Tbx1* probe detects mutated RNA also in the mutant (*Tbx1^lacZ^*^/lacZ^) functionally null. Arrows indicate regions of the trunk mesenchyme with *Meox1* and *Tbx1* signals overlap. Ph: pharyngeal cavity; Nt: neural tube; OFT outflow tract; IFT: inflow tract.

The scale bar is 100µm.

**Supplementary Figure 8**

RNA-scope images of whole-mount E9.5 embryos (right side of the embryo) (a-b) and sagittal sections (c-c’) using *Hoxb3* as the probe (red). In a-a’ is also shown the *Tbx1* probe signal (green). PA1, 2, 3: position of the pharyngeal arches 1, 2, 3, respectively. PPd: peripharyngeal expression domain of *Hoxb3*; ED: ectopic expression domain.

The scale bar is 100µm.

**Supplementary Figure 9**

Coronal sections of E9.5 mouse embryos (*WT and Tbx1^-/-^*) hybridized by RNAscope with *Tbx1* (green) and *Hoxb3* (red) probes. Arrows indicate the position of the *Hoxb3* ectopic expression domain in the mutant, posteriorly and partially surrounding the otocyst (dashed circles). Anterior is up, posterior is down.

The scale bar is 100µm.

