## Supplementary fugures for "Regulatory mutants of the *Tbx1* gene alter transcription programs of lineage determination and patterning in early mesoderm"

### Supplementary Figure 1

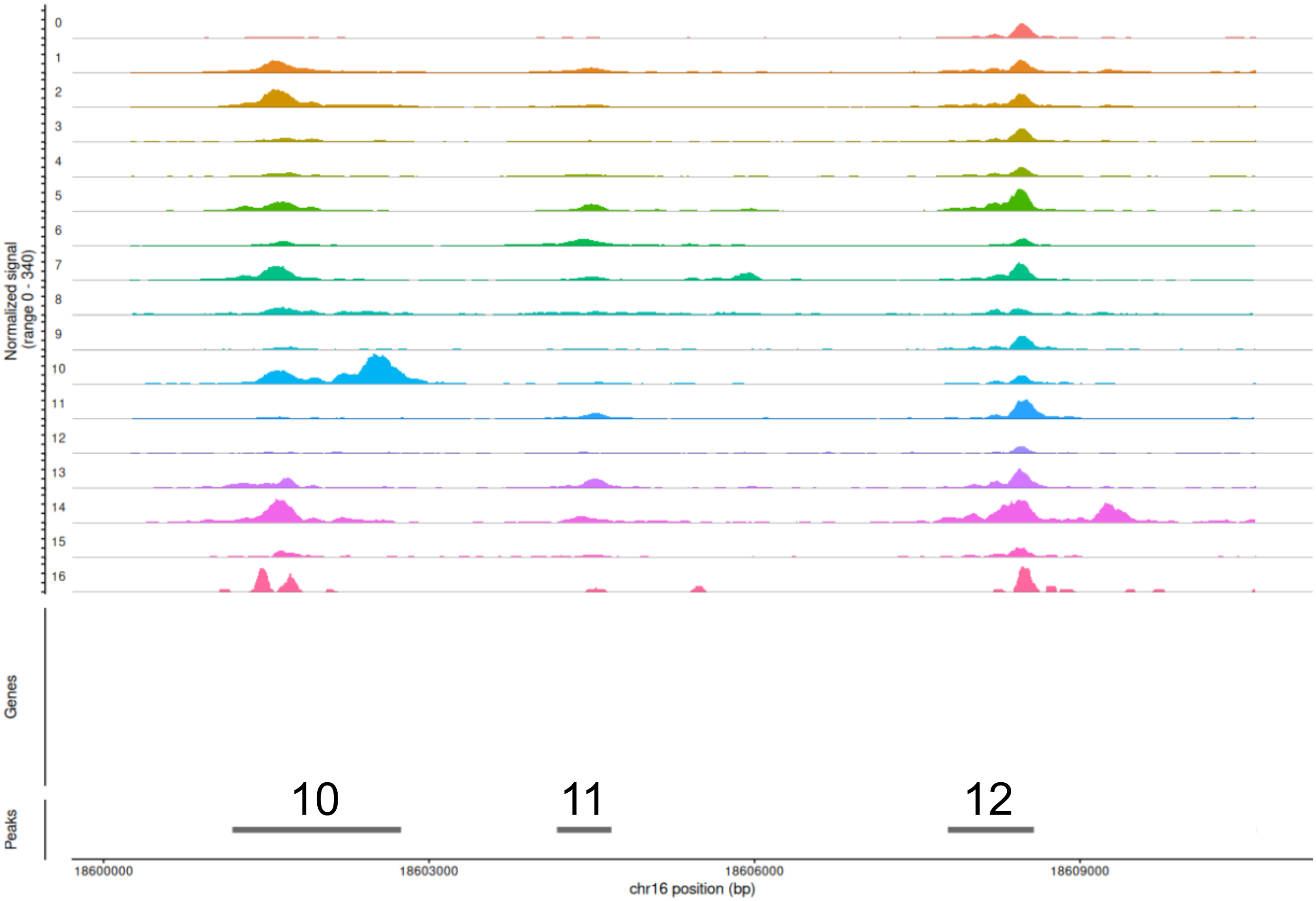

### Supplementary Figure 2

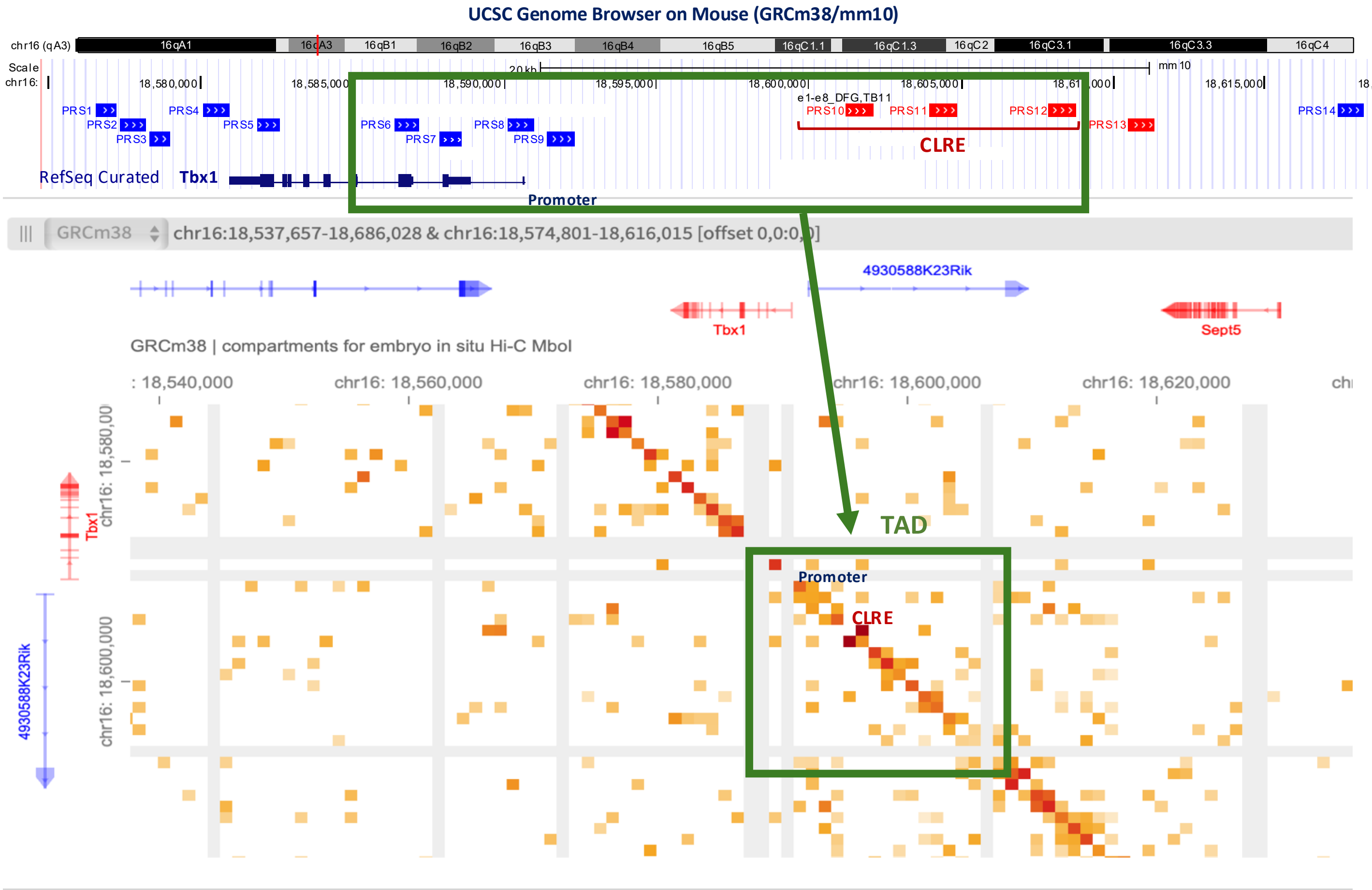

Supplementary Figure 3

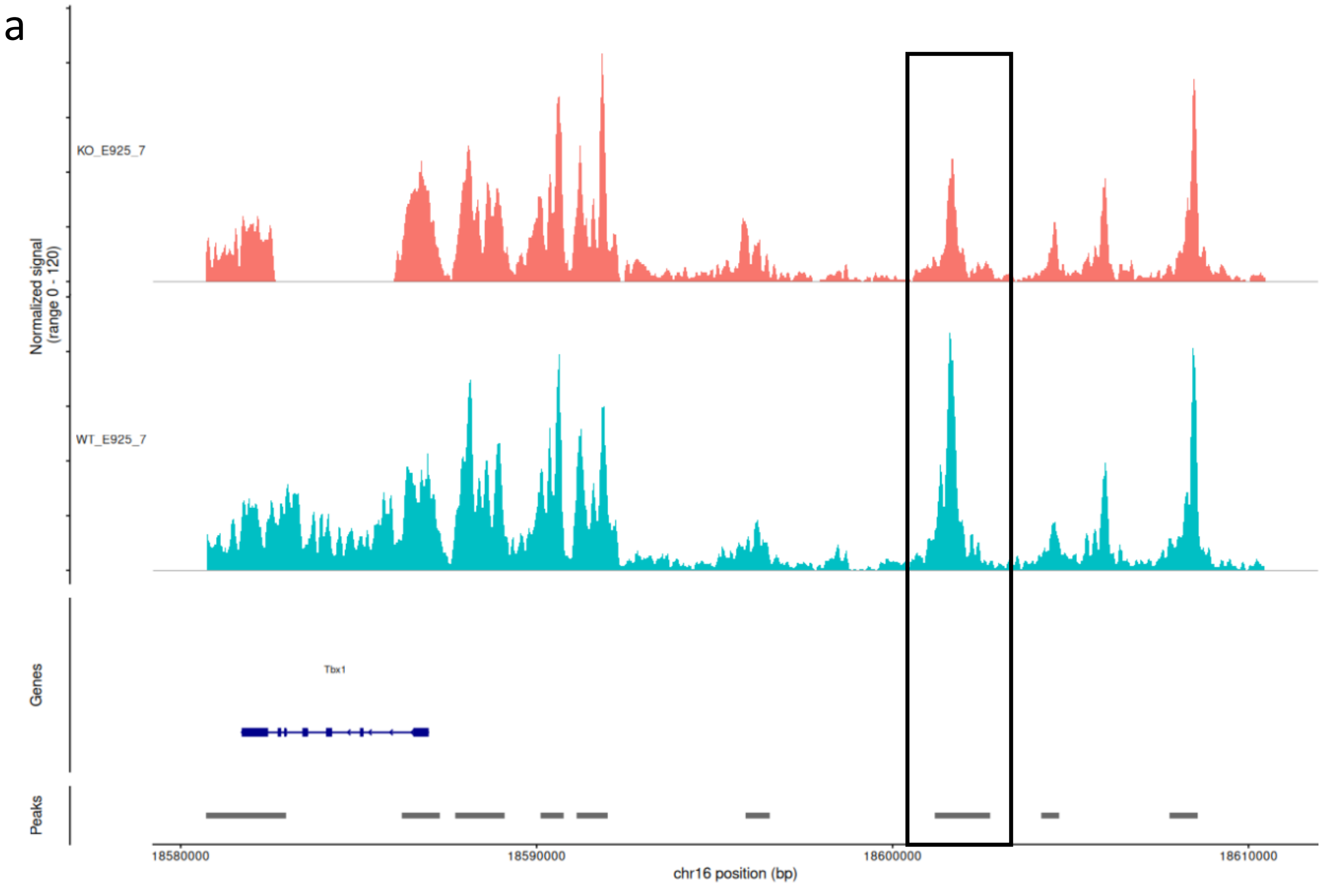


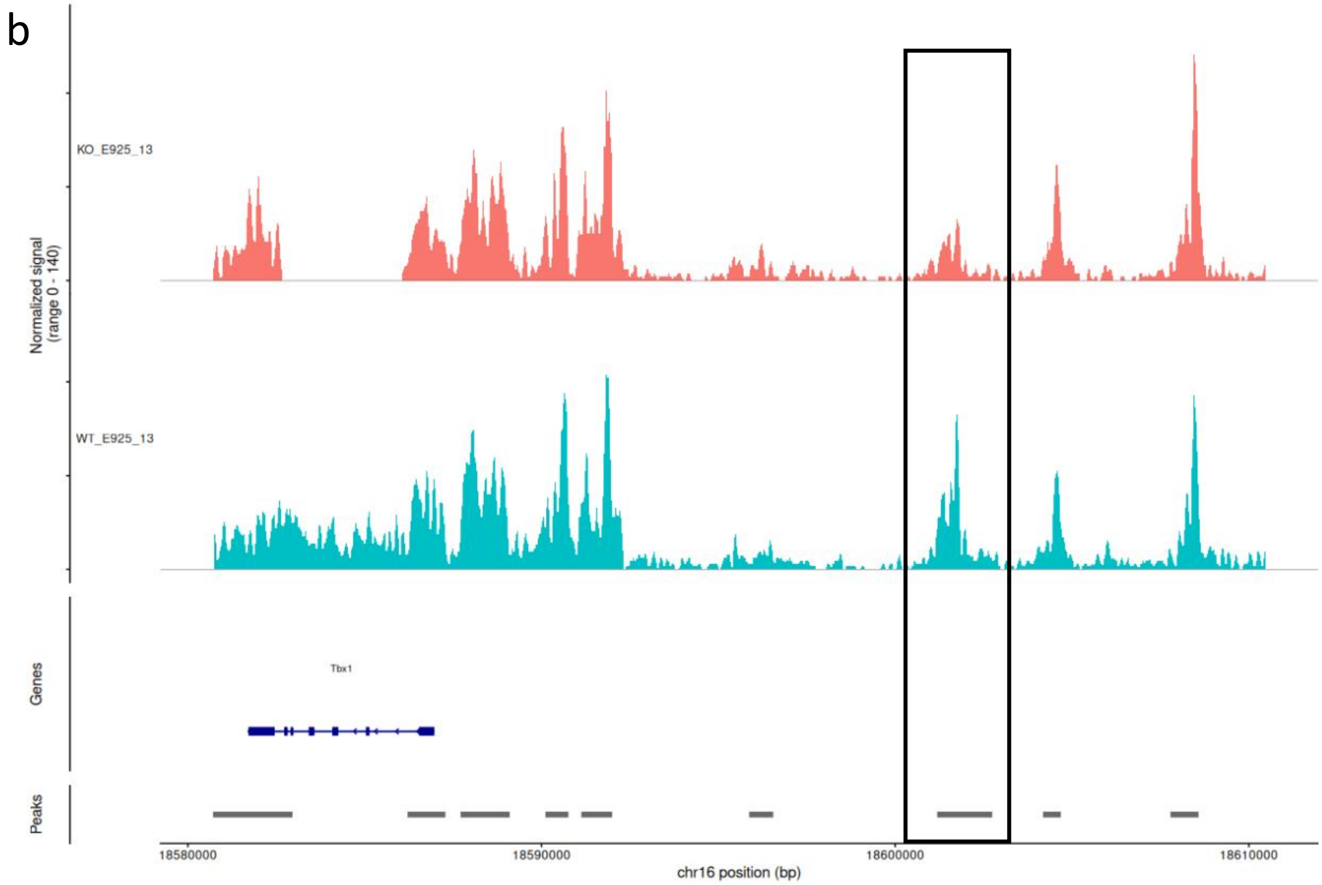

### Supplementary Figure 4

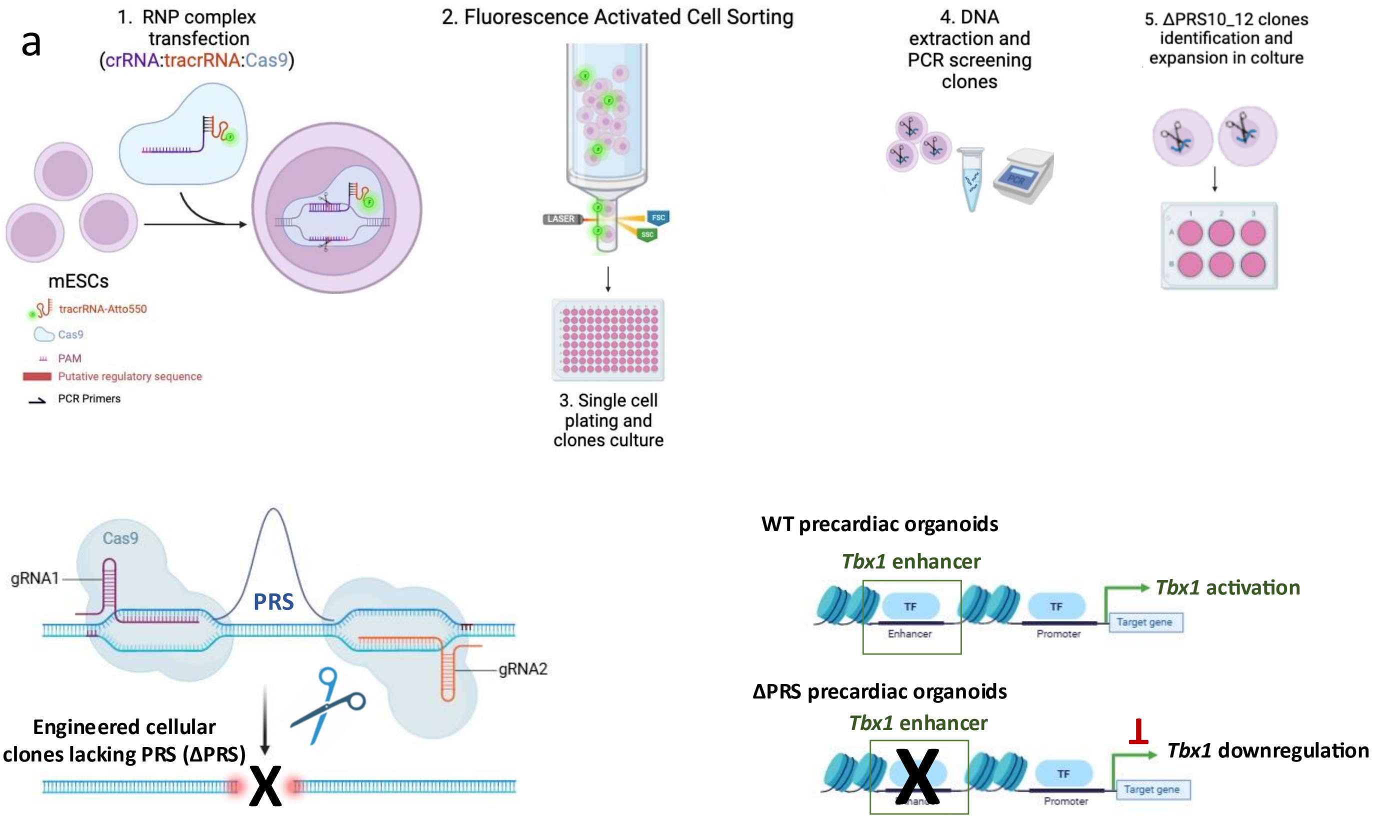


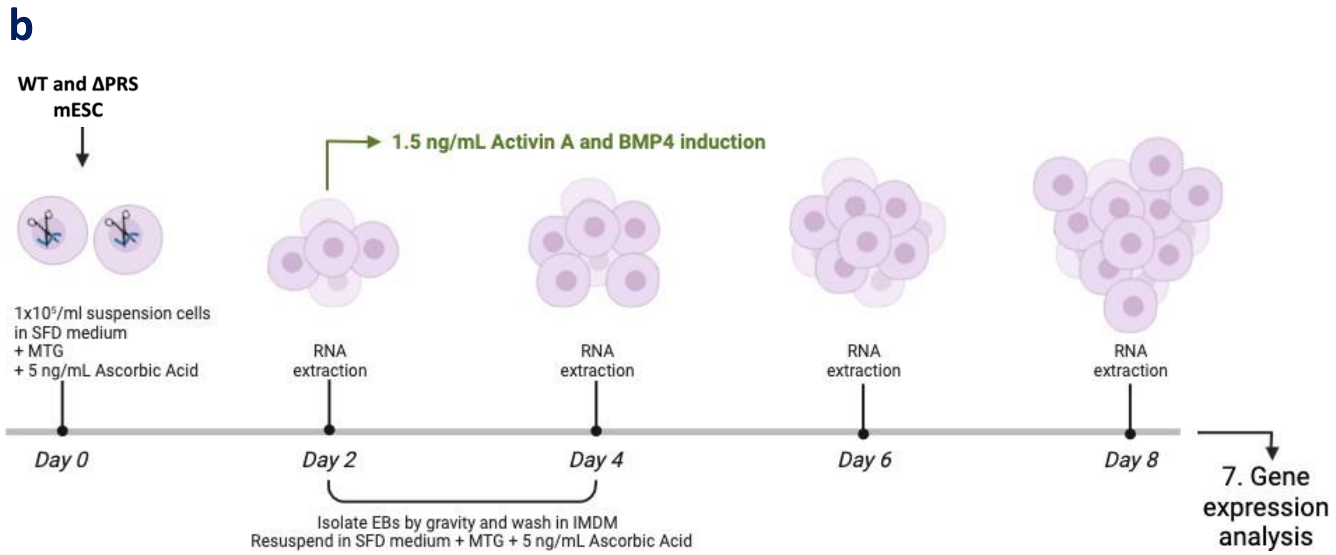

### Supplementary Figure 5

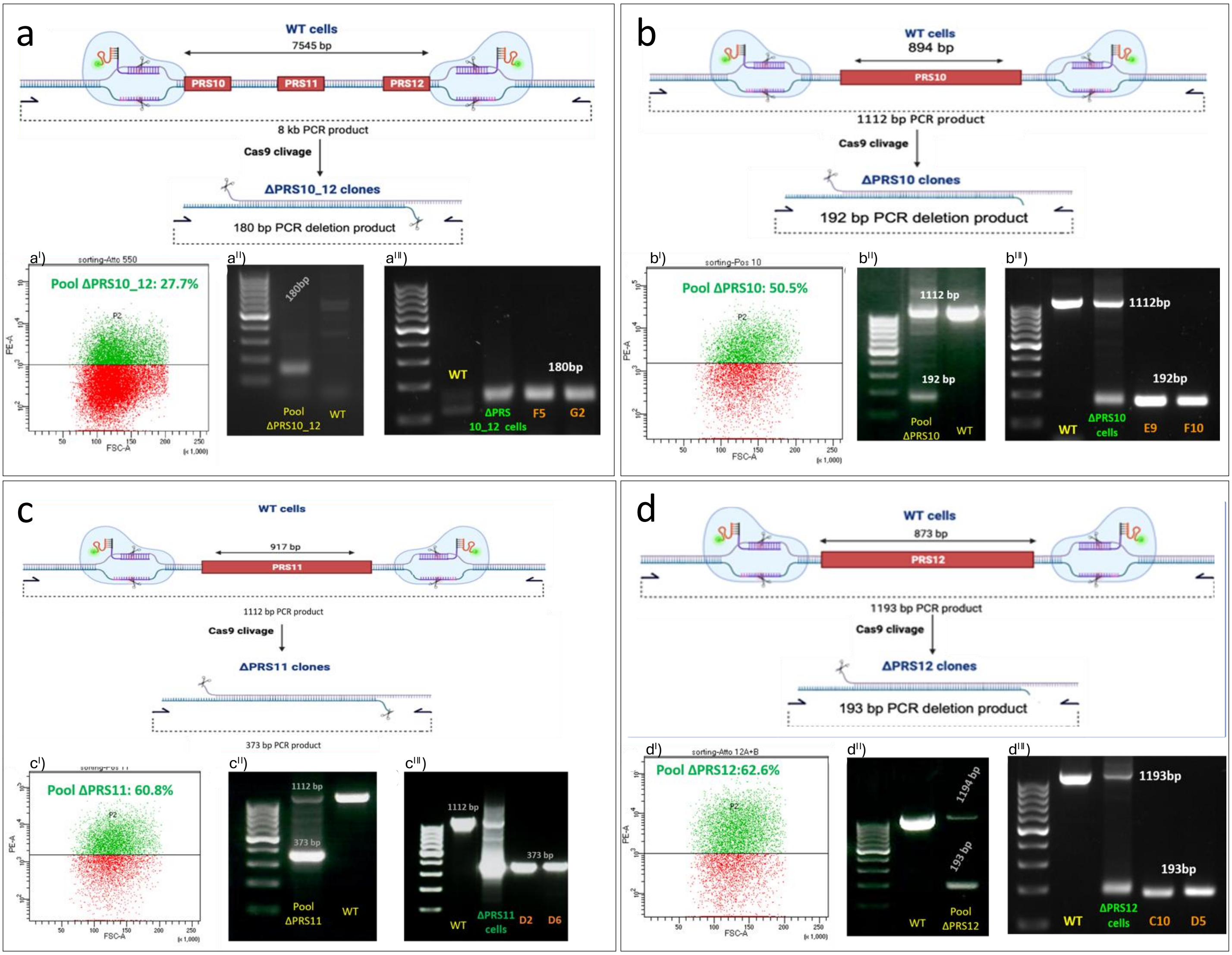

### Supplementary Figure 6

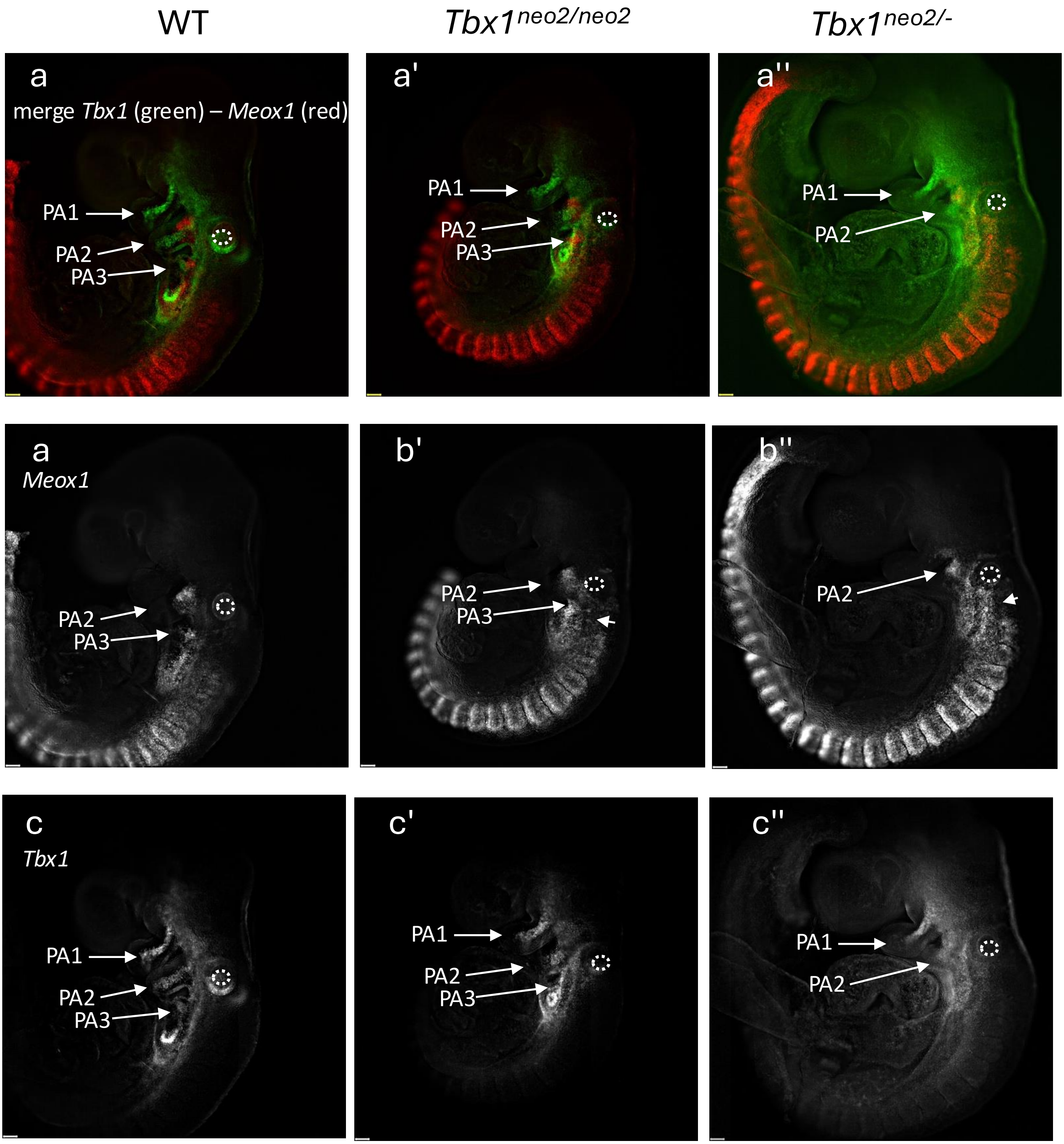

Supplementary Figure 7

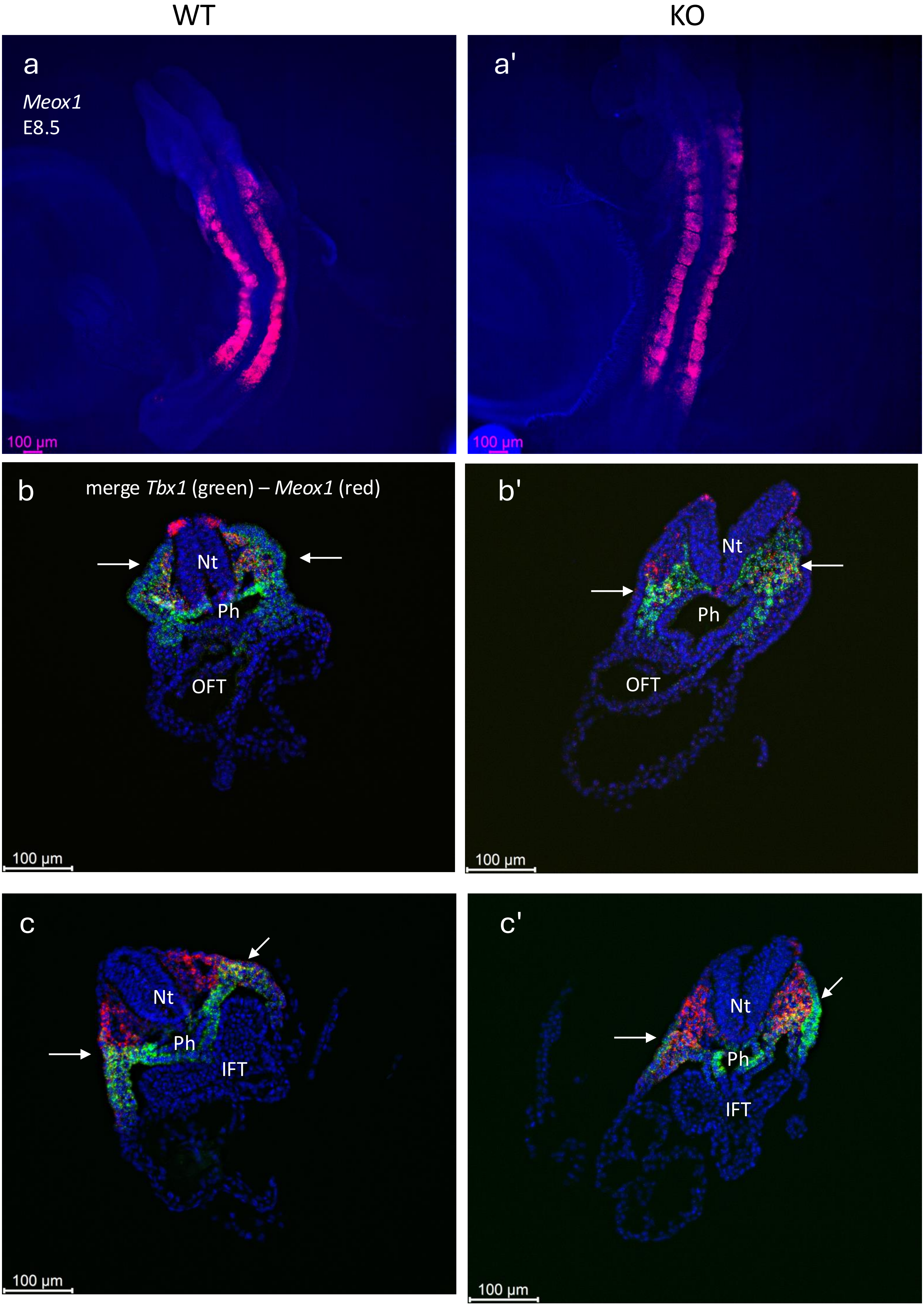

Supplementary Figure 8

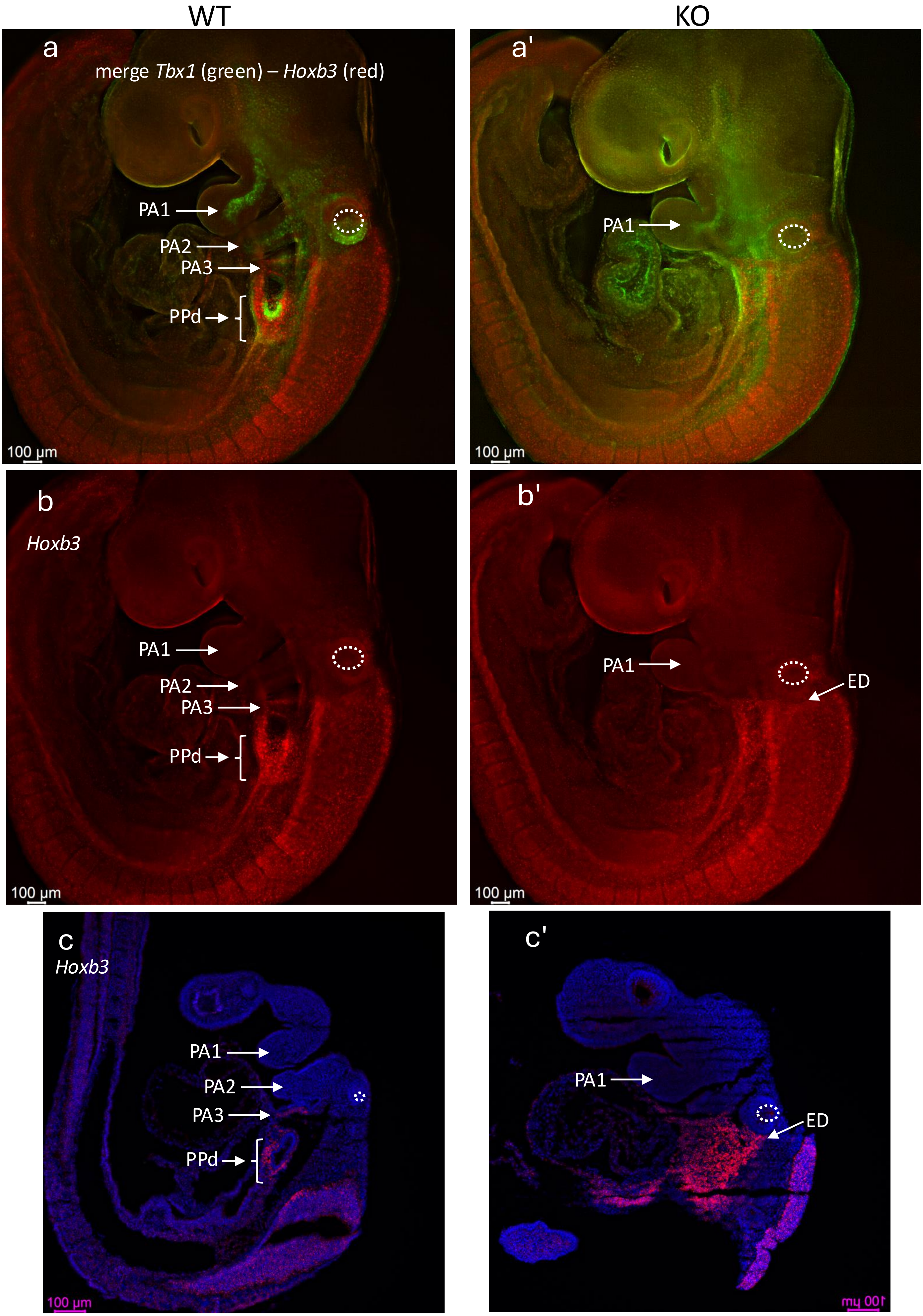

Supplementary Figure 9

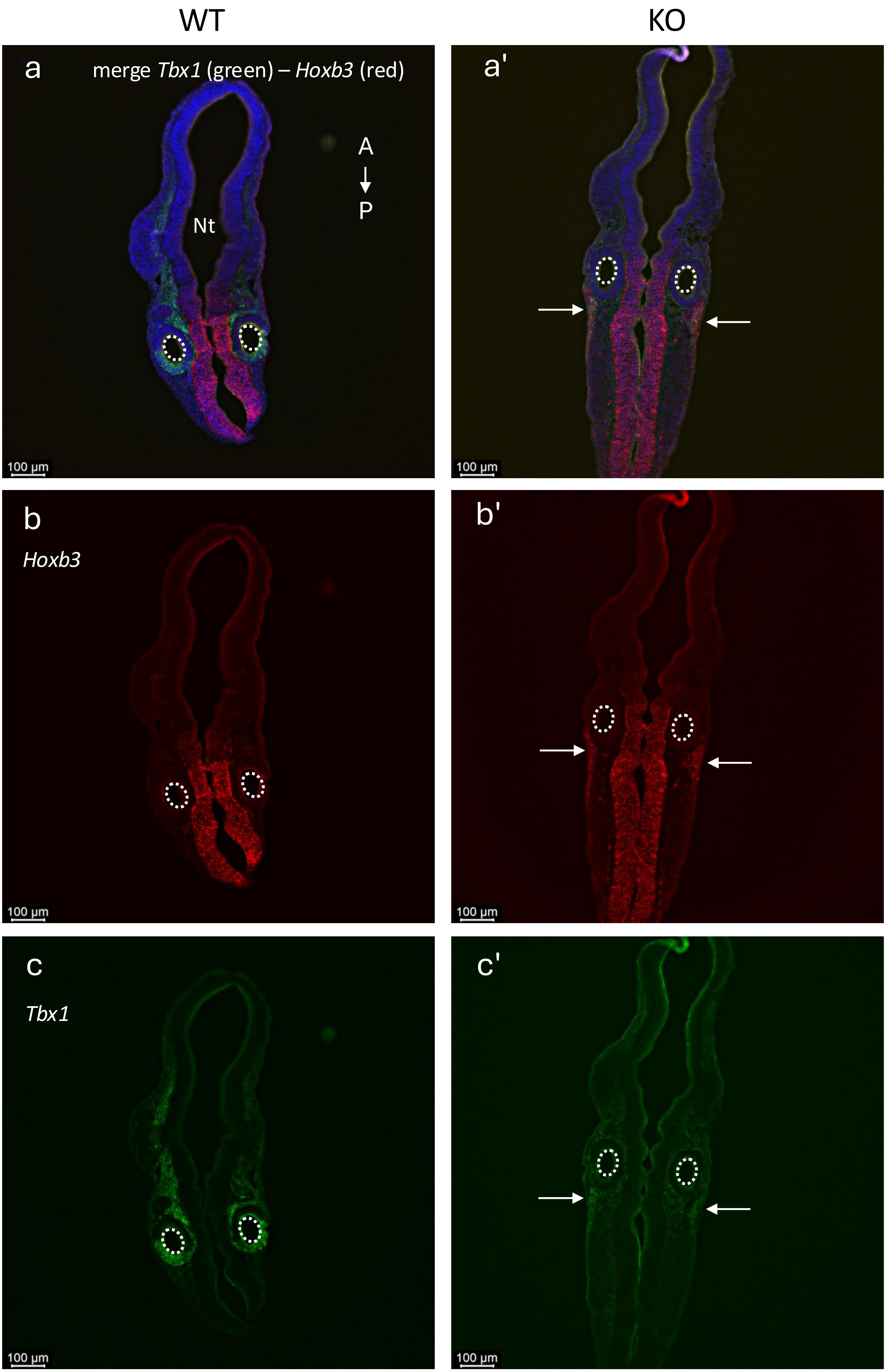
